# Gigapixel behavioral and neural activity imaging with a novel multi-camera array microscope

**DOI:** 10.1101/2021.10.05.461695

**Authors:** Eric E Thomson, Mark Harfouche, Pavan C Konda, Catherine Seitz, Kanghyun Kim, Colin Cooke, Shiqi Xu, Whitney S Jacobs, Robin Blazing, Yang Chen, Sunanda Sharma, Timothy W Dunn, Jaehee Park, Roarke Horstmeyer, Eva A Naumann

## Abstract

The dynamics of living organisms are organized across many spatial scales. However, current cost-effective imaging systems can measure only a subset of these scales at once. We have created a scalable multi-camera array microscope (MCAM) that enables comprehensive high-resolution recording from multiple spatial scales simultaneously, ranging from cellular structures to large-group behavioral dynamics. By collecting data from up to 96 cameras, we computationally generate gigapixel-scale images and movies with a field of view over hundreds of square centimeters at 5um sensitivity. This allows us to observe the behavior and fine anatomical features of numerous freely moving model organisms on multiple spatial scales, including larval zebrafish, fruit flies, nematodes, carpenter ants, and slime mold. The MCAM architecture allows stereoscopic tracking of the z-position of organisms using the overlapping field of view from adjacent cameras. Further, we demonstrate the ability to acquire dual color fluorescence video of multiple freely moving zebrafish, recording neural activity via ratiometric calcium imaging. Overall, the MCAM provides a powerful platform for investigating cellular and behavioral processes across a wide range of spatial scales by removing the bottlenecks imposed by single-camera image acquisition systems.

## Introduction

Complex biological systems typically include processes that span many levels of organization across spatial scales that vary by multiple orders of magnitude (Churchland and Sejnowski, 1988; Iain D. Couzin and Jens Krause, 2003). For instance, processes such as schooling (Wright and Krause, 2006) or shoaling (Larsch and Baier, 2018) in fish involve social interactions between multiple individuals across tens of centimeters, but also include coordinated changes in eye movements (Harpaz et al., 2021) and attendant fluctuations in neural and synaptic activity (Fan et al., 2019) at the micrometer scale. Currently, no single imaging system allows for unrestricted access to each scale simultaneously, which requires the ability to jointly observe a very large field of view (FOV) while maintaining a high spatial resolution. Hence, imaging systems typically make tradeoffs in their measurement process. For instance, existing neuronal imaging methods typically require an animal to be tethered (Ahrens et al., 2013; Harvey and Svoboda, 2007), which restricts the natural behavioral repertoire (Hamel et al., 2015) and inhibits spontaneous movement (Kim et al., 2017; O’Malley et al., 2004). Alternatively, closed-loop mechanical tracking microscopes (Johnson et al., 2020; Kim et al., 2017; Marques et al., 2020; Nguyen et al., 2016a; Reiter et al., 2018; Symvoulidis et al., 2017) have recently been developed in an attempt to address this challenge, but can only follow *single* organisms (Cong et al., 2017) and thus miss inter-organism interactions. Scanning-based imaging approaches are routinely employed to observe larger areas at high-resolution, either by mechanical movement (Potsaid et al., 2005), the use of microlens arrays (Orth and Crozier, 2013, 2012) or via illumination steering as a function of time (Ashraf et al., 2021; Gustafsson, 2005; Mudry et al., 2012; Zheng et al., 2014). Unfortunately, sequential imaging strategies are not able to *simultaneously* observe living processes at high resolution over a large area, such as movement of multiple individuals or even large collections of freely moving organisms, which will naturally change position during the slow acquisition cycle, thus generating ambiguities.

To simultaneously observe multiple organisms over a large area at high spatial resolution, one appealing idea is to increase the FOV of a standard microscope by producing a single large-diameter, large numerical aperture lens that captures images at the diffraction limit across the full FOV of interest. Unfortunately, geometrical optical aberrations increase with the surface area of a lens when its diameter increases (Lohmann, 1989). The number of corrective lenses required to compensate for these aberrations quickly increases, as does the overall lens system scale, in a nonlinear manner (Lohmann, 1989). These additional corrective lenses rapidly increase the size, weight, complexity and cost of imaging optics (Brady et al., 2018; Mait et al., 2018) which generally limits most commercially available microscope lenses to transfer less than 50 megapixels of resolved optical information to the image plane (Zheng et al., 2014). Complex lens systems specifically designed to address this challenge have not been able to increase this quantity beyond several hundred megapixels (McConnell et al., 2016; McConnell and Amos, 2018; Sofroniew et al., 2016). At the same time, the largest digital image sensors currently available also contain at most several hundred megapixels (“Canon LI8020SA 250MP CMOS Sensor,” 2021), which suggests a natural limit to single lens imaging systems precluding direct acquisition of gigapixel-scale image data. Recently, a 3.2-gigapixel digital camera for a specialized Large Synoptic Survey Telescope was developed, but its cost, $168 million, is prohibitive for most lab-based research.

To overcome these challenges, and extend observational capabilities to the gigapixel regime, we designed and constructed an inexpensive, scalable *Multi Camera Array Microscope* (MCAM) (**Figure 1A**). The MCAM collects data in parallel from an arbitrarily sized rectangular grid of closely spaced modern CMOS image sensors (each with ~10^7^ pixels, that are 1 μm in size, **Supplementary Figure 1**), which are currently produced in large volumes for the cell phone camera market and are thus readily available in many varieties and at low-cost. Each MCAM image sensor is outfitted with its own high-resolution lens to capture image data from a unique sample plane area (**Figure 1A**, zoom in). The MCAM sensors and lenses are arranged such that their fields-of-view partially overlap with their immediate neighbors (**Supplementary Figure 2**), which ensures that image data are jointly acquired from a continuous area (**Figure 1B**). Such a parallelized image acquisition geometry (Brady et al., 2018) can, in theory, be scaled to produce an arbitrarily large FOV. Here, we present MCAM designs with up to 96 cameras that allow us to record information-rich bright-field gigapixel videos of multiple freely moving species, including Carpenter ants (*Camponotus* spp.), fruit flies (*Drosophila melanogaster*), nematodes (*Caenorhabditis elegans*), zebrafish (*Danio rerio*) and the amoeboid slime mold (*Physarum polycephalum)*(Nakagaki et al., 2000) at 18 μm full-pitch optical resolution (i.e., the ability to resolve two 9 μm bars spaced 9 μm apart) over a 384 cm^2^ FOV. To help distill these large image sets into salient and actionable data, we adapted a variety of modern image analysis tools, including a convolutional neural network-based object tracker, to work at gigapixel scale.

**Figure 1.**
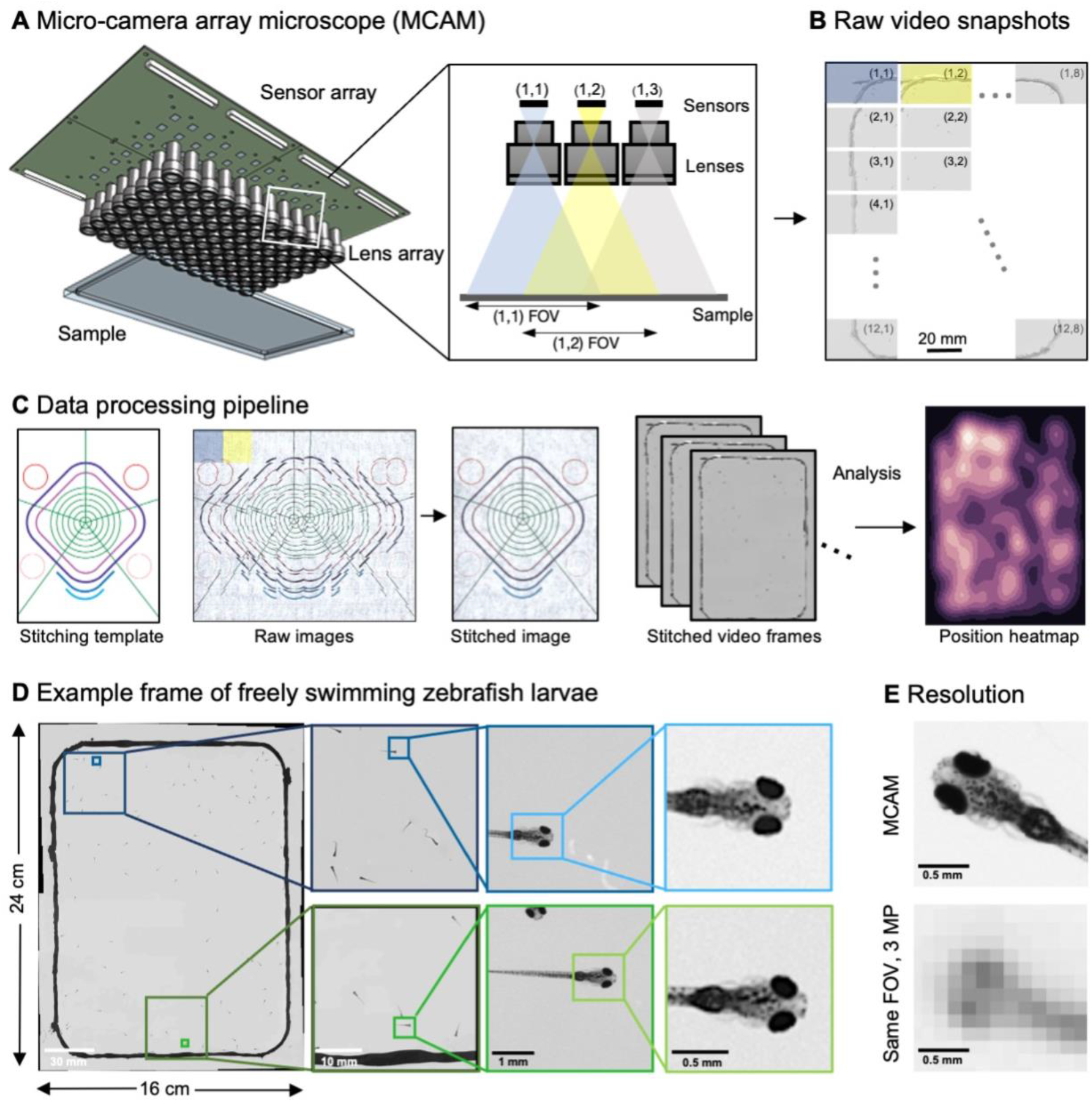
Architecture of the multiple camera array microscope. **A.** Schematic of MCAM setup to capture 0.96 gigapixels of snapshot image data with 96 micro-cameras. Inset shows MCAM working principle, where cameras are arranged in a tight configuration with some overlap (10% along one dimension and 55% along the other) so that each camera images a unique part of the sample and overlapping areas can be used for seamless stitching and more advanced functionality. **B.** Example set of 96 raw MCAM frames (8 x 12 array) containing image data with an approximate 18 μm two-point resolution across a 16 x 24 cm field-of-view. **C.** Snapshot frame sets are acquired over time and are combined together via a calibration stitching template to reconstruct final images and video for subsequent analysis. An example of an analysis is a probability heatmap of finding a fish within the arena at a given location across an example imaging session. **D.** Example of one stitched gigapixel video frame of 93 wildtype zebrafish, 8 days post fertilization, selected at random from an hour-long gigapixel video. **E.** Resolution comparison of an example larval zebrafish captured by the MCAM and from a single-lens system covering a similar 16 x 24 cm FOV using a 3 MP image sensor (but capable of high-speed video capture).

The MCAM technology also enables a new suite of imaging experiments for monitoring fluorescent neuronal activity in *populations* of freely moving animals. After outfitting the imaging system for epi-fluorescence imaging, we visualized fluorescent neurons in Drosophila larvae and recorded neural activity in multiple freely swimming zebrafish expressing neuronal genetically encoded calcium indicators such as GCaMP6 (Dana et al., 2016). Ratiometric analysis of dual color recordings of neuronal activity across many simultaneously freely moving zebrafish revealed response patterns related to specific locomotor behaviors. As a unique approach to address the FOV/resolution tradeoff, the MCAM offers a scalable platform that overcomes current imaging bottlenecks and allows us to simultaneously observe phenomena spanning from the microscopic to the macroscopic regimes.

## Results

### Multi Camera Array Microscope (MCAM) design and image processing

To achieve large-FOV microscopic imaging, we first designed and constructed a modular MCAM imaging unit that consisted of a 4 × 6 rectangular array of 24 individual Complementary metal–oxide–semiconductor (CMOS) image sensors (10 megapixels per sensor, 1.4 μm pixels) and associated lenses (25 mm focal length, 0.03 numerical aperture, 12 cm^2^ FOV per lens). We integrated the 24 sensors onto a single circuit board (**Supplementary Figure 1**), and directly routed all data through a custom-designed control board containing a set of field-programmable gate arrays (FPGAs). The FPGAs deliver the image data to a single desktop computer via a USB3 cable, where the raw high-resolution video from all 24 sensors is saved for subsequent post-processing. The total pixel count per snapshot from each 24-camera array is 0.24 gigapixels.

To verify that our modular MCAM architecture could scale to image acquisition from much larger regions, we joined four such 24-sensor units together in a two-by-two arrangement into a *gigapixel imaging system* (**Figure 1A, Supplementary Movie 1**), which streamed video data from 96 cameras simultaneously (0.96 gigapixels total raw data; **Supplementary Figure 1E**) to a single desktop computer. We successfully operated this final system at full pixel count at up to 1Hz frame rate, yet implementation of on-sensor pixel binning and/or improved data transmission arrangements can yield higher frame rates (see Discussion). A continuous recording session at 0.74 Hz lasted almost one hour resulted in 2.5 TB of unstitched data (2649 frames, 962MB per frame) and 1.17 TB stitched data (442 MB per image). The system exploits the rapid current read/write speeds for individual solid state drives on a single personal computer (achieving approx. 1GB/s), and marks a significant speedup from previously published gigapixel imaging systems (Brady et al., 2012).

To assess the MCAM resolution, we imaged a custom-designed resolution test target extending across the 96-camera MCAM’s entire 16 cm x 24 cm FOV (**Supplementary Figure 2**) to demonstrate an 18 μm full-pitch optical resolution cutoff, which corresponds to a maximum spatial frequency cutoff of 147 line pairs/mm under incoherent illumination (Ou et al., 2015). We additionally demonstrate the single-point sensitivity (i.e., the ability to reliably detect isolated features of specific size) of the MCAM to fall between 3-5 μm, by imaging a mixed set of sparsely arranged 3 μm and 5 μm diameter polystyrene microspheres and confirming detection with a high-resolution (20x) objective lens (**Supplementary Figure 2**). Importantly, we emphasize that the above resolution and sensitivity performance is independent of the number of included cameras within the MCAM, thus providing a high-resolution imaging platform that can scale to an arbitrary FOV when more cameras are included in the array.

Processing vast amounts of data -- sometimes millions of images -- from multiple cameras presents many unique opportunities and challenges compared to standard single-camera imaging systems. For all data, we first saved raw images to disk followed by a two-step offline preprocessing procedure. We performed standard flat-field correction to compensate for vignetting and other brightness variations within and between cameras (**Methods**). For each MCAM snapshot in time, we then digitally stitched the images acquired by all of the micro-cameras together into a single composite image frame (**Figure 1C, Supplementary Figure 3, Methods**). Note that while stitching is not technically required for many subsequent image analysis tasks, it is often useful for visual presentation of acquired results, such as organism trajectory assessment or multi-organism location measurement (e.g., heatmap in **Figure 1C**). Following preprocessing, images then entered various application-specific machine learning-driven algorithms, as outlined below.

### High-resolution, large-FOV imaging of large groups of larval zebrafish

After validating the hardware and implementing our basic image acquisition workflow, we used custom Python code to produce gigapixel videos from the image data streamed directly to four separate hard drives. To record the behavior of ~100 wildtype, 8 days post-fertilization (dpf) zebrafish (**Supplementary Movie 2**), we placed them into a 22 × 11 cm 3D-printed (resin on glass) behavioral arena (**Figure 1D, Methods**). As each gigapixel image frame contains information about the zebrafish at multiple spatial scales, from the social structure of the population down to fine-grained morphological features of individual fish, we were able to resolve details like pectoral fin position and the distribution and morphology of individual melanophores, i.e., the pigmented skin cells on the body, in addition to arena-wide swim trajectories (**Figure 1D**). In an MCAM image, a zebrafish (8 dpf) is captured by 9699 ± 2758 pixels (n = 93 zebrafish), which is significantly higher than in standard imaging systems. For example, when using a single 20 MP camera with the same large-FOV, fish would be represented by less than 400 pixels, and even less in older juvenile fish (Romero-Ferrero et al., 2019). For a standard 3 MP, a comparable FOV a fish occupies less than ~100 pixels (**Figure 1E**).

To automatically detect and crop individual organisms for subsequent analysis, we adapted a convolutional neural network (CNN)-based object detector to accommodate our stitched MCAM image frames (**Figure 2A**; **Supplementary Figure 5; Methods**), which are otherwise too large for training or inference on existing graphical processing units (GPUs). Using a custom data augmentation pipeline (**Methods**), a tailored transfer learning approach (Sarandi et al., 2018) yielded a network that could accurately detect individual zebrafish even when occluding one another (**Supplementary Figure 5I, Supplementary Movie 3**).

**Figure 2.**
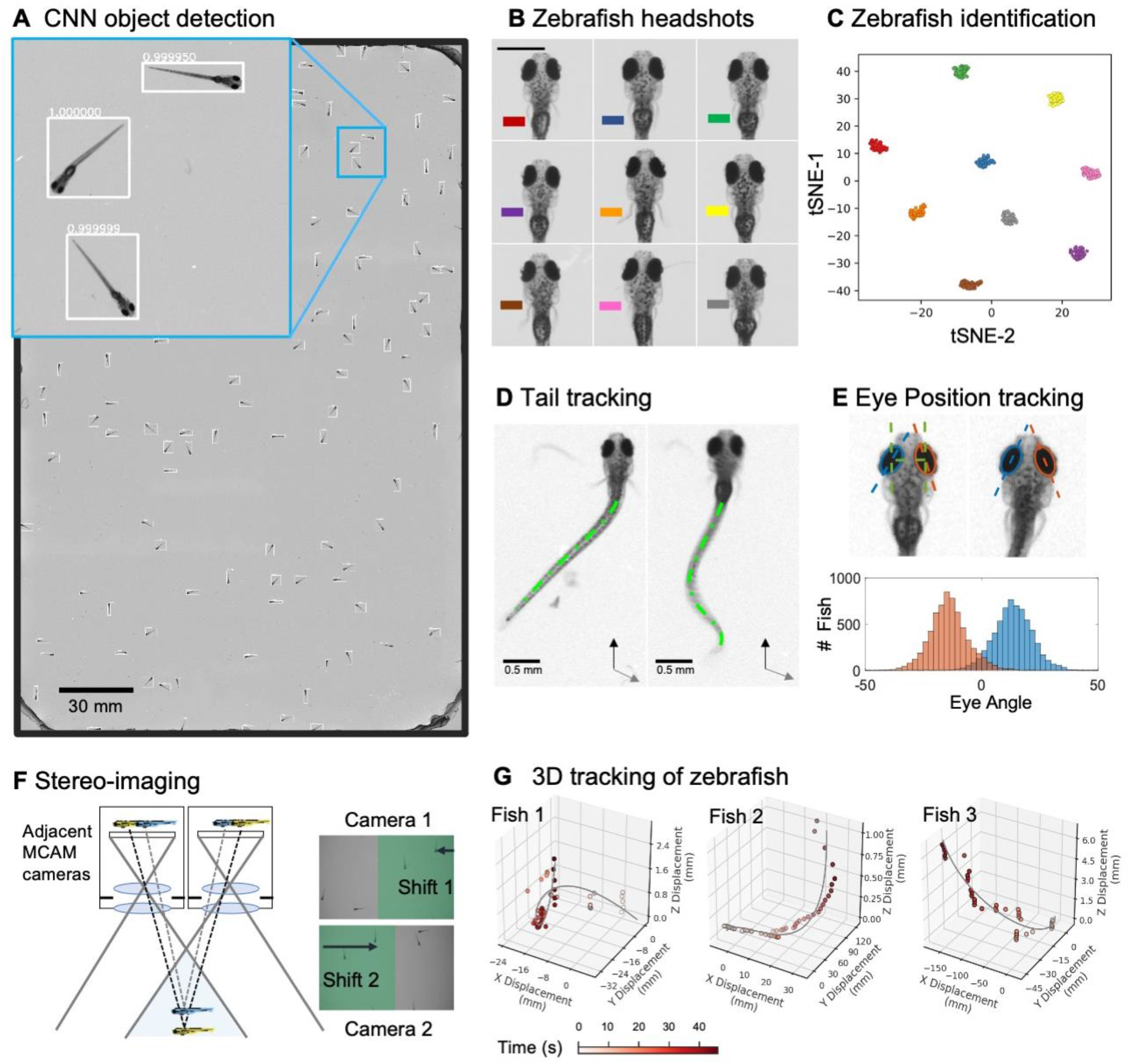
Gigapixel high-resolution imaging of collective zebrafish behavior. **A.** Example frame of stitched gigapixel video of 130 wildtype zebrafish, 8 days old. An object detection pipeline (Faster-RCNN) is used to detect and draw bounding boxes around each organism. Bounding boxes include detection confidence score (for more details see Methods). Each bounding box’ image coordinates is saved and displayed on stitched full frame image. Zoom in shows three individual fish (blue box). Scale bar: 1cm. **B.** Headshots of nine zebrafish whose images used to train a siamese neural network (see Methods, Supplementary Figure 6). Colors indicate the fish identity used to visualize performance in **C**. Each fish has a distinct melanophore pattern, which are visible due to the high resolution of MCAM. Scale bar: 1 mm. **C.** Two-dimensional t-SNE visualization of the 64-dimensional image embeddings output from the siamese neural network, showing that the network can differentiate individual zebrafish. For the 9 zebrafish, 9 clusters are apparent, with each cluster exclusively comprising images of one of the 9 fish in **B**, suggesting the network is capable of consistently distinguishing larval zebrafish. **D.** Close-up of three zebrafish with tail tracking; original head orientation shown in gray (bottom right). **E.** Automated eye tracking from cropped zebrafish (see methods for details); results histogram of eye angles measured on 130 zebrafish across 20 frames. **F.** Optical schematic of stereoscopic depth tracking. The depth tracking algorithm first uses features to match an object between cameras (top) and then calculates the binocular disparity between the two cameras to estimate axial position with an approximate resolution of 100 μm along z. **G.** 3D trajectory of 3 example larvae from recorded gigapixel video of 130 wildtype zebrafish, with z displacement estimated by stereoscopic depth tracking.

After automatically detecting zebrafish in each MCAM frame, analysis was performed on image data extracted from within each bounding box containing an individual fish, vastly reducing memory and data handling requirements (**Figure 2A**, inset). To track individual zebrafish over time, we exploited the high spatial resolution of the MCAM to consistently recognize individuals by morphological differences (**Figure 2B**), which has not been shown before in zebrafish younger than 21 days old. Specifically, we trained a siamese neural network (Bertinetto et al., 2016) (**Supplementary Figure 6**) that consistently grouped regions of interest containing the same individual zebrafish together (**Figure 2C**). Given the high spatial resolution of the MCAM, this network had unique access to idiosyncrasies in melanophore patterns, structural and overall body morphology when learning to distinguish individual animals. In addition, we used the MCAM’s resolution to identify behavioral details of each zebrafish by performing tail tracking (**Figure 2D**) and eye position measurement (**Figure 2E**), which yielded aggregate statistics over populations of zebrafish during free swimming.

In addition, the MCAM’s multi-lens architecture offers novel measurement capabilities as compared to standard single-lens microscopes. One example is stereoscopic depth tracking. By using rectangular image sensors (16:9 aspect ratio), the MCAM provides images from adjacent cameras that overlap approximately 50% in the x-direction and <10% in the y-direction to facilitate stitching (**Supplementary Figure 2**). This overlap ensures that any point in the sample plane is simultaneously observed by at least two cameras and allows us to use positional disparities from matched features (i.e., stereoscopic depth measurement techniques) to calculate the depth of objects within adjacent camera image pairs (**Figure 2F**, **Methods**). Briefly, after matching features within adjacent frames using a scale-invariant feature transform, pixel-level lateral feature displacements generate depth estimates using known MCAM imaging system parameters (e.g., inter-camera spacing and working distance). Using this stereoscopic depth measurement approach yields approximately 100 μm accuracy in depth localization over a >5 mm depth range (**Supplementary Figure 7**). To demonstrate this stereoscopic capability of the MCAM, we tracked the 3D trajectory of zebrafish within the entire swim arena (**Figure 2G**). Example 3D tracking revealed typical quantifiable signatures of fish spontaneous swim trajectories along spatial axes typically inaccessible (Bolton et al., 2019) without dual view or camera systems.

### Using MCAM for simultaneous behavior and fluorescence imaging

The high-resolution and large field of view make the MCAM a powerful tool for recording cellular-level detail in freely moving organisms. To examine this, we integrated fluorescence imaging capabilities in a separate 24-sensor MCAM unit (**Figure 3A**). Specifically, we added two sets of custom excitation sources (high-powered light emitting diodes (LEDs)) to illuminate the entire arena and mounted a bank of 24 emission filters via a 3D printed filter array holder over each camera lens (**Methods**; **Supplementary Figure 8A,B**). With these two simple additions, the MCAM easily captured epi-illuminated fluorescence images from each MCAM sensor (**Supplementary Figure 8C,D**). We imaged freely moving third instar drosophila larvae (**Figure 3B**), expressing GCaMP in the brain and ventral nerve cord (Grueber et al., 2003; Vaadia et al., 2019). The epi-fluorescent setup allowed us to observe an extremely bright convergence of proprioceptive neurons in the anterior/ventral regions of the larva (**Supplementary Movie 11**). Similarly, **Figure 3C** shows GFP expression localized to the cranial nerves of a 6 day old zebrafish *Tg(isl1a:GFP)* (Higashijima et al., 2000).

**Figure 3.**
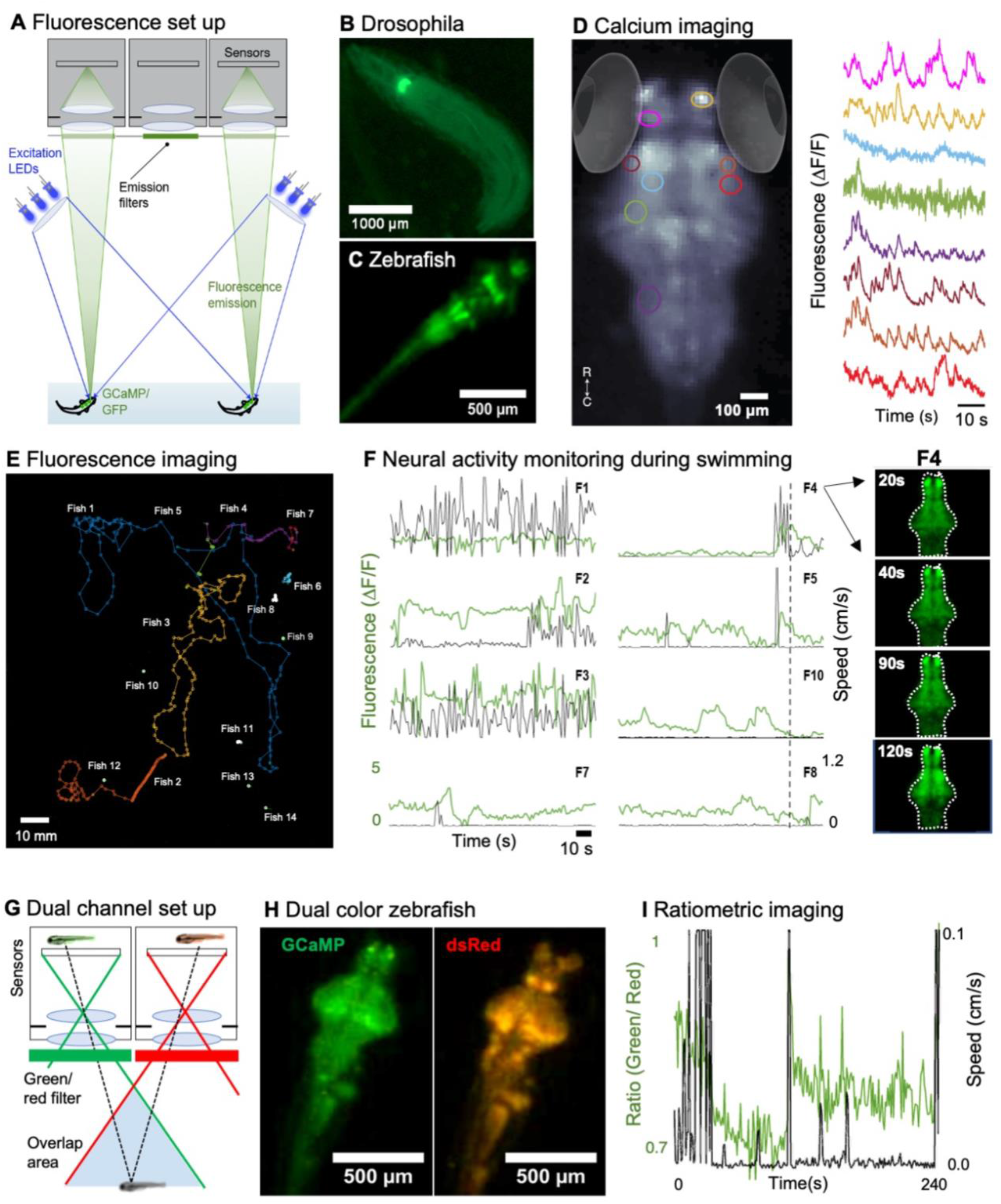
Fluorescence imaging of neural activity with MCAM. **A.** Schematic of MCAM fluorescence imaging set up. **B.** Example fluorescent MCAM image zoom of fluorescent D. melanogaster instar 3 larva (410Gal4 x 20XUAS-IVS-GCaMP6f), showing fluorescence can be detected in proprioceptive neurons. See Supplementary Movie 11. **C.** Localized expression of GFP in cranial nerves and brain stem motor neurons in *Tg(Islet1:GFP)* transgenic zebrafish. **D.** Average fluorescent image of agarose embedded zebrafish with pan-neuronal expression profile of *Tg(elavl3:GCaMP6s)*. Colored selections represent region of interests (ROI) shown to the right. Average ΔF/F traces of these ROIs show that differences in neural activity are easily resolved by the MCAM. See Supplementary Movie 4. **E.** Fourteen freely moving zebrafish, *Tg(elavl3:GCaMP6s)* in a 24-camera MCAM system, tracked over a 120s video sequence. **F.** Fluorescent neural activity (ΔF/F) of six example zebrafish from **E**. over 120s plotted with swim speed shows heightened activity during locomotion for some fish. While changes are relatively constant activity during continuous swimming (e.g., F1), signals for fish with limited mobility can be clearly observed (e.g., F7). **G.** Dual channel fluorescence imaging configuration. As each area is imaged by two MCAM sensors, simultaneous recording of red and green fluorescence permit ratiometric imaging. **H.** Example dual-channel images of the same double transgenic zebrafish *TgBAC(slc17a6b:LOXP-DsRed-LOXP-GFP)* x *Tg(elavl3:GCaMP6s)*. **I**. Ratiometric analysis of pan-neuronal neural activity for an example zebrafish during freely swimming plotted as swim speed. Fluorescence increases are visible right after certain movement events.

We monitored neural activity by acquiring fluorescence images in the “single-sensor” mode (10 frames/sec.) from zebrafish expressing the genetically encoded fluorescent calcium indicator GCaMP6s (Chen et al., 2013) across almost all neurons across the brain via the *elavl3* promoter (**Figure 3D**). To reduce motion artifacts, we started by embedding zebrafish in low melting point agarose and recorded spontaneous neural activity for 5 min (**Supplementary Movie 4**).The ΔF/F traces over time in different brain regions of interest (ROIs) in **Figure 3D** clearly show region-specific neuronal activation profiles.

Next, we combined fluorescence and behavioral tracking, using 24-camera MCAM to record video at 1 frames/sec. for 5-minute imaging sessions of 14 freely moving zebrafish within an 8×12 cm swim arena (**Figure 3E**). **Figure 3F** shows the fluorescent activity in six of these fish by tracking, centering, and aligning the brains of each fish (**Figure 3F**, *right*). For zebrafish that move sporadically clear increase and decrease of activity can be detected after the onset of swimming, and a marked quenching of activity when swim bouts ceased. While the MCAM in the current configuration cannot detect single-neuron fluorescence activity in the zebrafish brain, the data do show that our multi-sensor architecture can measure regionally localized functional brain activity data in freely moving fish. Calibrated videos of freely swimming transgenic zebrafish with pan-neuronal GCaMP6s expression verify that our system can non-invasively measure neural activity in >10 organisms simultaneously during natural interactions (**Figure 3E-F**). As signals from moving fish will always represent the challenge to distinguish calcium increases from movement artifact, we utilized the MCAM’s FOV overlap to perform ratiometric imaging. To improve the accuracy of fluorescence measurement during free movement, we installed alternating red and green fluorescent filters to image both red and green fluorescence (**Supplementary Figure 8E**), enabling us to perform ratiometric imaging(Nguyen et al., 2016b). Here, we imaged double transgenic zebrafish *Tg(slc17a6b:LOXP-DsRed-LOXP-GFP-)* x *Tg(elavl3:GCaMP6s)* (Dana et al., 2019; Koyama et al., 2011) which express a red fluorescent protein (DsRed) in GABAergic neurons and the green calcium sensor in almost all neurons. Since the red fluorescent protein is not changing, the red channel image serves to normalize the GCaMP6s fluorescence and therefore can compensate for movement artifacts that might otherwise be interpreted as neural activity dependent fluorescence changes (**Supplementary Figure 8F,G**). In **Supplementary Figure 8** and **Supplementary Movie 5** we show that we can track and measure neural activity in two separate zebrafish while ratiometrically recording their respective fluorescent brain activity (**Figure 3G-H**). We determined (**Figure 3I**) that whole brain calcium traces increase during times of increased locomotor activity, similar as found in drosophila (Maimon et al., 2010) and mice (Dombeck et al., 2007; Niell and Stryker, 2010).

### Application to multiple model organisms

In this section, we expand beyond the zebrafish, showing the value of the MCAM as a general, scalable imaging platform for the study of multiple model systems in biology. **Figure 4** demonstrates gigapixel recording of nematode (*Caenorhabditis elegans*) behavior in three different strains: wild type (N2), an unpublished transgenic line (NYL2629) that expresses GFP and mCherry in motor neurons, and a mutant (unc-3) line (Brenner, 1974) that has a loss-of function mutation in the unc-3 gene (unc-3 is expressed in motor neurons and is activated when worms become hungry (Prasad et al., 1998)). We plated these worms on a large, square (24.5cmx24.5cm) petri dish and employed a CNN-based segmentation algorithm (**Supplementary Methods**). We measurements indicate ~58,000 worms per frame, captured at about 450 pixels per worm within a typical gigapixel video (averaged across 20 frames, single frame in **Figure 4C**). Surprisingly, both the unc-3 mutants, and especially the NYL2629 strain show previously unreported swarming behavior. Approximately 3 days after being transferred to a new nutrient lawn, the NYL2629 strain began to form tight swarms that tiled most of the petri dish in a semi-periodic fashion (**Figure 4B**), as has been observed in other strains (de Bono and Bargmann, 1998). Based on the hypothesis that this could be driven by partial loss of function in the reporter strain, we then tested for such effects more directly in an unc-3 mutant, where we observed qualitatively different swarming behavior: large wavefronts of worm aggregates (2-3 cm wide) (**Figure 4C**). Notably, such aggregative behavior was not observed in the wildtype line (**Figure 4A**). Using computer vision algorithms to track worms and detect swarm formation shows how such large-scale differences in behavior can be observed with the MCAM (**Supplementary Movies 6-8**). Recent research on collective behavior has demonstrated that swarming behaviors are modulated by various factors, including oxygen levels and animal density (Demir et al., 2020; Ding et al., 2019). While further investigation of the underlying causes and spatiotemporal features of swarming in these different strains is needed, our data clearly demonstrate that the MCAM is a useful tool for the quantitative study of factors that control social clustering and swarming in C. elegans (**Figure 4E**).

**Figure 4.**
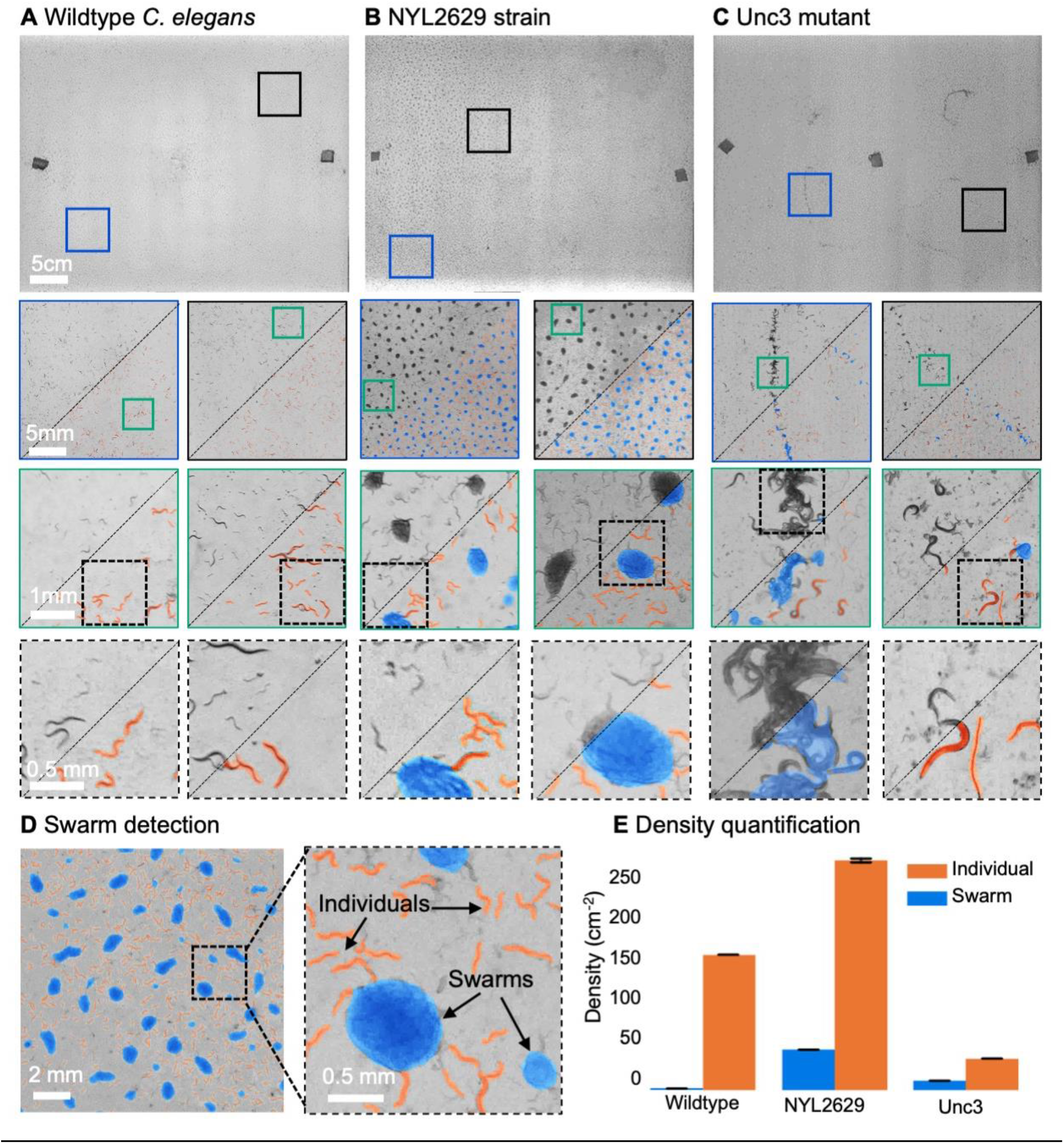
MCAM imaging of swarming behavior in C. elegans. **A.** Wild type C. elegans (N2 line) spread across the entire arena uniformly. Consecutive zoom ins show no significant swarming behavior. **B.** NYL2629 C elegans strain shows a periodic tiling of the petri dish over the majority of the arena as C. elegans form swarm aggregates that can be automatically detected. **C.** Unc3 mutant C. elegans exhibit large wavefront swarms of activity, without the periodic tiling seen in the NYL2629 strain. Unc3+ genotype inhibits a significant swarming behavior and causes swarms, annotated in blue, all over the entire arena. Our system provides sufficient resolution across a large field-of-view to enable studying behavioral patterns of C. elegans. In all cases the imaging was performed after day 3 of starting the culture. Supplementary Movies 6-8 allow viewing of full video. Entire gigapixel video frames viewable online. **D.** Segmentation of swarms (blue) and individual C. elegans (orange) using a U-shaped convolutional network allows the automatic creation of binary identification masks of both individual worms and swarms. After segmentation, we used a pixel-connectivity-based method to count the number of objects (worms or swarms) in each gigapixel segmentation mask. **E.** Bar graph quantifying density of worms within and outside of swarms for the different strains in A-C. Error bars indicate S.E.M for 5 cm sizes samples across the arena.

Another species that displays stark behavioral and morphological variations on spatial scales spanning multiple orders of magnitude is the slime mold *Physarum polycephalum* (Alim et al., 2013). This single-celled, multinucleate protist exhibits complex behavior at large spatial scales. For instance, its pseudopodia will envelope nutrients in its environment, foraging over a search path spanning many hundreds of square centimeters (Gawlitta et al., 1980; Rodiek and Hauser, 2015). Exploiting this, we observed the well-known ability of the slime-mold to traverse mazes, though on much larger spatial scales (**Figure 5A; Supplementary Movie 10**), larger than previously observed (Nakagaki et al., 2000). Also, using time-lapse imaging with a 24-camera MCAM array (0.24 gigapixels), we acquired data over a 46-hour period, showing the cumulative growth and exploration of a standard 10 cm diameter petri dish that had been seeded with oatmeal flakes (**Supplementary Movie 13**). At the same time we observed the well-known cytoplasmic flow within individual plasmodia (Durham and Ridgway, 1976), involving particulate matter tens of square microns, flowing across multiple centimeters, which we were able to observe in individual MCAM cameras (**Supplementary Movie 14**).

**Figure 5.**
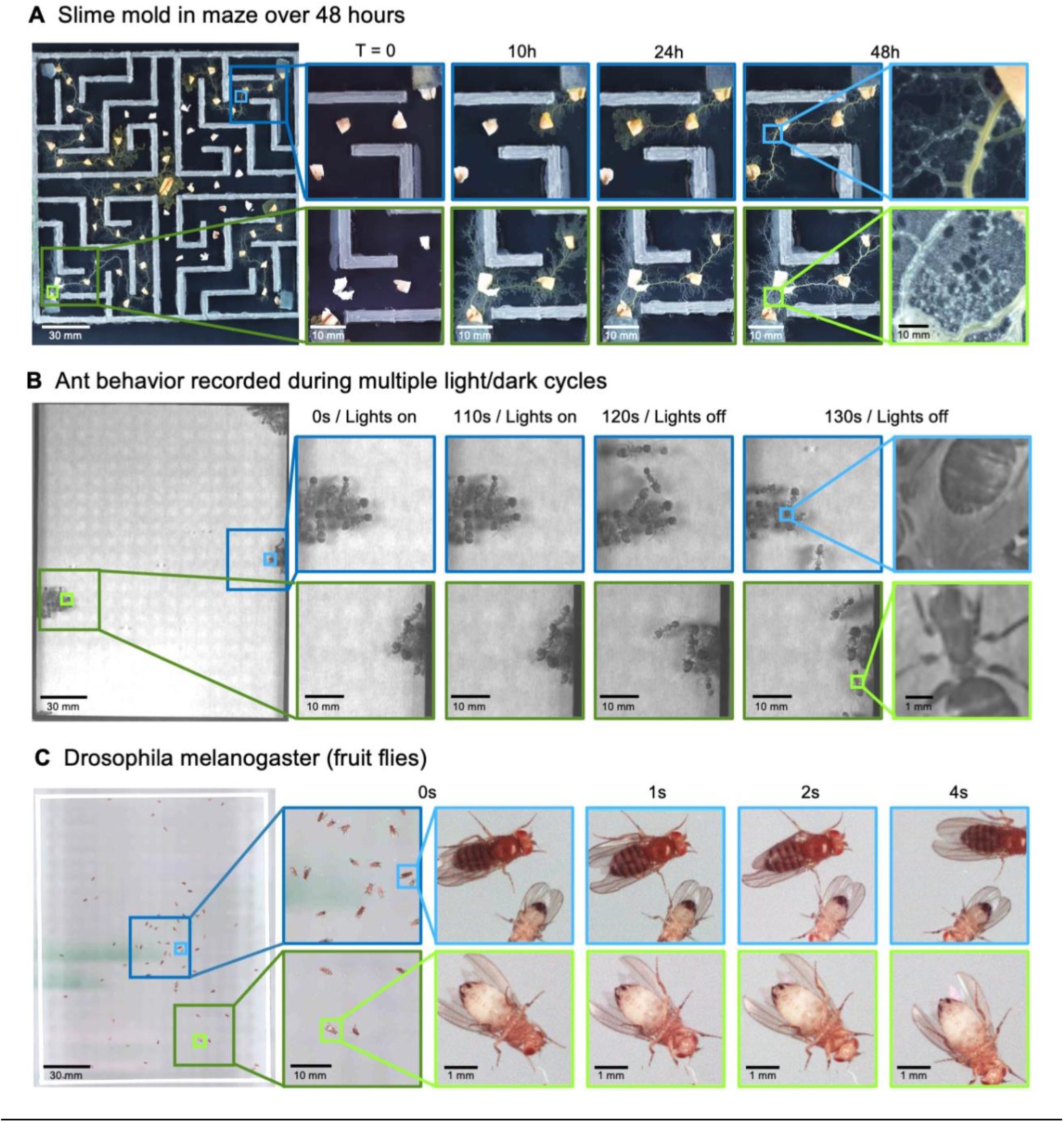
MCAM imaging of behaviors in multiple small organisms. **A.** Gigapixel video of slime mold (*P. polycephalum*) growth on maze (16.5 cm per side). Four mold samples were taken from a seed inoculation on another petri dish and placed in four corners of the maze. Recording for 96 hours shows the slime mold grow within the maze. Supplementary Movies 10, 13, and 14 highlight the MCAM’s ability to simultaneously observe the macroscopic maze structure while maintaining spatial and temporal resolution sufficient to observe microscopic cytoplasmic flow. **B.** Gigapixel video of a small colony of Carpenter ants during light and dark alternations. Overall activity (velocity) of ants measured with optical flow increased during light off versus light on phases. Full MCAM frame observes Carpenter ant colony movement and clustering within large arena, while zoom-in demonstrates spatial resolution sufficient for leg movement and resolving hair on abdomen. See Supplementary Movie 9. **C.** Adult Drosophila gigapixel video frame showing collective behavior of several dozen organisms at high spatial resolution, with insets revealing fine detail (e.g., wings, legs, eyes) during interaction. See Supplementary Movie 12.

Ant colonies show a highly distributed multi-scale spatial structure, with many individuals spending time in their base colony, and the foragers leaving the colony to retrieve food (Mersch et al., 2013). To demonstrate how the MCAM could be useful to study such behavior, we imaged behavior in the Carpenter ant (*Camponotus pennsylvanicus*)(Hamilton et al., 2011). We observed dramatic changes in behavioral arousal with change in light levels. Carpenter ants are nocturnal animals, and their activity levels are very tightly locked to ambient light (Narendra et al., 2017; Sharma et al., 2004). When we alternated light and dark cycles with small groups of ants, we observed clear fluctuations in activity levels, including the dissolution of tight clusters of “sleeping ants” with increased movement (**Figure 5B**), but also social interactions and grooming (pulling on antennae with legs) (**Supplementary Movie 9**).

Finally, the fruit fly, *Drosophila melanogaster* has become a benchmark model system for the genetic analysis of behavior(Datta et al., 2008), development (Tolwinski, 2017), and neurophysiology (Mauss et al., 2015). We recorded fly behavior at multiple developmental stages using the MCAM. As discussed above, we imaged behavior and neuronal GFP expression in third-instar larvae. Furthermore, the MCAM allowed us to observe large-scale social behavioral patterns in adult fruit flies, such as face-to-face interactions, and mutations in the wings such as curling (**Figure 5C, Supplementary Movie 12**). At the same time, we could see small features of individual flies, such as hairs on individual legs, grooming behavior, and the sex of each animal (**Figure 5C**, inset). This demonstrates the potential use for the MCAM in large-scale genetic screening in addition to the study of collective behavior that involves interactions on very fine spatial scales.

## Discussion

We report on the development of a multi-camera array microscope (MCAM), constructed from closely tiling multiple individually inexpensive, powerful image sensors, to overcome the limitations of single-objective and single-sensor systems, allowing the acquisition of unprecedented high-resolution images over a large, in principle infinite FOV. While prior designs have considered multiple microscopes to image distinct regions in parallel (Weinstein et al., 2004), or via cascaded multiscale lens designs (Fan et al., 2019) that can yield light-field-type imaging measurements (Broxton et al., 2013; Levoy et al., 2009), the MCAM provides a novel, flexible arrangement that can monitor phenomena in freely moving organisms at multiple spatial scales. The number of cameras can be varied to produce different FOVs with varying frame rates, depending on the targeted imaging application. The working distance can also be changed to jointly increase or decrease the degree of inter-camera FOV overlap to unlock novel functionality. By adopting a parallelized approach to video acquisition, the MCAM removes a longstanding bottleneck of current single-camera microscope designs to provide researchers with greater flexibility, in particular for observing model organisms at high-resolution in an unconstrained manner.

We have also demonstrated two key capabilities of the MCAM’s multi-lens architecture – 3D tracking and dual-channel fluorescence imaging. The former functionality opens up a new dimension to behavioral analysis, while the latter can increase both the information content and accuracy of quantitative fluorescence measurement in moving specimens. We demonstrated how joint acquisition of green, fluorescent calcium indicators and red fluorescent proteins yields a ratiometric measurement of neuronal activity during free movement of zebrafish, but there are likely a wide variety of alternative exciting functionalities that MCAM overlapped sampling can facilitate.

While we were able to sample gigapixel images at 1Hz for up to 60 minutes – a significant increase in speed and duration from previous published efforts (Brady et al., 2012) – most biological applications naturally require faster acquisition rates. For instance, our current acquisition speed of 1 Hz is too slow to determine specific patterns in swim kinematics in freely swimming zebrafish (Marques et al., 2018), additional fast low resolution imaging from below could resolve this in the future (Johnson et al., 2020). Yet, the most straightforward paths to directly increasing the MCAM frame rate are 1) adopting an alternative data transfer scheme (e.g., replacing USB data transmission with PCIe), and 2) executing on-chip pixel binning, image cropping, and/or lossless compression to increase temporal sampling rates at the expense of spatial measurement count. Both improvements are currently under development in a re-designed MCAM system. A third promising direction is to dynamically pre-process acquired images on the included FPGAs before directing them to computer memory to lighten data transfer loads.

Such pre-processing could range from acquiring only from cameras that contain objects of interest to on-the-fly tracking and cropping of individual organisms or features of interest. We see the infrastructure embodied in the current MCAM implementation as an auspicious initial scaffolding upon which we can build such natural extensions to provide researchers with maximally salient image data spanning the microscopic and macroscopic regimes.

While this work primarily focused on examining how the MCAM can enhance model organism experiments, our new imaging technology can also be applied to a variety of other applications. Large area, high-resolution imaging is important in industrial inspection of manufactured goods (Yang et al., 2020) and materials (Ngan et al., 2011), as well as semiconductor metrology (Huang and Pan, 2015). For instance, such methods are required for defect detection on semiconductor wafer surfaces. It can also unlock new functionalities in imaging large pathology specimens, such as whole-mount brain slices (Axer et al., 2011). Biological researchers may also be able to utilize the MCAM platform to monitor bacterial colonies (Shi et al., 2017) and cell cultures over macroscopically large areas. Such a wide variety of potential applications provides many new directions to guide development of array imaging in the future.

In summary, our MCAM architecture fits within a growing body of computational imaging hardware (Mait et al., 2018) that is specifically designed not for direct human viewing, but instead with the assumption that data post-processing will be used to extract relevant information for subsequent analysis. While novel display technology currently enables visual assessment of the MCAM’s gigapixel videos that can be many Terabytes in raw size, we anticipate that simultaneously improving processing software (e.g., 3D organism tracking, machine learning-assisted analysis) will prove critical to fully realize the microscope’s benefits. As image sensor technology continues to improve with such automated processing moving on-chip (Amir et al., 2018; Posch et al., 2014), we anticipate that our parallelized imaging architecture can help overcome the standard tradeoff between resolution and field-of-view while at the same time yield manageable information-rich datasets to provide new insight into complex biological systems.

## Methods

### MCAM Electronics

Our first constructed gigapixel MCAM system consists of 96 individual complementary metal oxide semiconductor (CMOS) sensors, lenses, and associated read-out electronics that transmit recorded image data from all sensors to computer memory for offline processing. The sensors and lenses (i.e., micro-cameras) are arranged in an aggregate 8×12 array, comprising 4 individual 24-sensor MCAM imaging units placed in a 2×2 array, as sketched in **Supplementary Figure 1**. Each CMOS sensor (Omnivision, OV10823) contains 10 megapixels (4320 × 2432) and a pixel size of 1.4 um for a 6 mm x 3.4 mm active area. In the prototype images included here, each CMOS sensor collected color information using a Bayer mosaic of red, green, and blue filters that are arranged in a pattern collecting 50% green, 25% red and 25% blue light. The sensors were placed at a 19 mm pitch, and their rectangular active area allows us to capture overlapping field-of-view information primarily along one dimension for stereo-based depth tracking and dual-channel fluorescence imaging. The imaging system’s total number of recorded pixels per frame is 0.96 gigapixels. Several results presented in this paper were from a single MCAM imaging unit that contains only 24 individual micro-cameras in a 4×6 layout (same micro-camera pitch and design), producing 240 recorded megapixels per frame.

When operating in full-frame mode, each CMOS image sensor delivers image data to an associated field-programmable gate array (FPGA) via custom-designed electronic circuitry. 6 CMOS sensors are controlled by one FPGA. A total of 16 sensor-connected FPGAs thus controlled and routed our 96-camera system’s image data to the computer. Each of these 16 FPGAs routes data from the 6-sensor clusters to one of four USB3.0 lines. To control data transmission, a “main” FPGA assigns each group of 4 FPGAs to port data (from 24 micro-cameras total) over one USB3.0 line (**Supplementary Figure 5)**. The 4 USB3.0 lines, each routing data from 24 CMOS sensors, are connected to a single desktop computer (Dell, Intel(R) Xeon(R) W-2133 3.6 GHz CPU), then saves directly to a set of 1TB solid-state hard drive memory cards. Each solid state drive with peak write speeds of more than 1GB/s (3x Samsung 970 Prom and 1x Samsung 960 Pro). We note that current solid-state memory drives offer a powerful means to store terabytes of data at rates of multi-gigabyte per second and are a primary driving technology for enabling collection of this extremely large-scale video data on a single desktop computer.

The MCAM offers two modes of operation. First, it can operate in a “full-frame” mode that records video data from all 96 micro-cameras at an aggregate frame rate of approximately 1 frame per second (see timing diagram in **Supplementary Figure 4B-C**). The per-frame exposure time for each micro-camera can be reduced to approximately 1 ms to limit any per-frame motion blur. While we are able to collect data from each CMOS sensor at a much higher frame rates, our current electronic design offers a limited data transmission bandwidth to computer memory when in “full frame” mode, which we address by FPGA-routing one image frame per second from each of the 96 micro-cameras and ignoring the remaining captured frames pre-transmission. Such transmission bandwidth limitations have been resolved in our next generation of MCAM (see Discussion).

The MCAM architecture also offers the flexibility to read data from a smaller number of sensors at a higher frame rate. For example, a “single-sensor” mode of operation allows us to selectively record video data from up to 4 electronically assigned micro-cameras at a frame rate of approximately 12 frames per second (**Supplementary Figure 5**). By allowing rapid evaluation of image quality during sample placement and focus adjustment, “single-sensor” mode is critical during device setup and can lead to dynamic spatiotemporal data acquisition schemes (e.g., fully digital organism tracking at 12 fps) in future designs. Current design efforts are focused on significantly improving frame rates both in full-frame and single-sensor mode to enable electronic tracking of rapid organism movement.

### MCAM Optics

Each of the 96 cameras utilizes a multi-element 12 mm diameter mount lens (Edmund Optics 58-207, 25 mm focal length, 2.5 f-number), with an 18 mm maximum outer diameter that currently sets the MCAM’s inter-micro-camera spacing (to 19 mm). As shown in **Supplementary Figure 2**, these lenses offer approximately 18 μm full-pitch resolution at a 150 mm working distance across an image FOV that extends across the entire image sensor (6 mm width). These lenses can also provide a bright-field sensitivity of at least 5 μm, confirmed by imaging with an alternative digital microscope using a 20x objective lens (0.5 NA, see **Supplementary Figure 2**).

We selected a working distance of approximately 150 mm to ensure that the images recorded by adjacent micro-cameras overlap sufficiently to enable seamless image stitching (see Methods, Software: Image Stitching). By using rectangular image sensors with a height/width ratio of 1/1.8, the current MCAM design offers approximately 52% image overlap along the longer sensor dimension when the lenses are configured to provide approximately 10% image overlap along the shorter image dimension. This overlapping field-of-view configuration is diagrammed in **Supplementary Figures 2 and 4** and guarantees that any point within the object plane (neglecting edge regions) is imaged by at least two micro-cameras. As noted above, this overlapped imaging configuration enables us to track the depth of various objects within the scene using stereovision-based depth tracking algorithms (see **Methods**, Software: Depth detection), as well as obtain dual-channel fluorescence measurements.

### MCAM Wide-field illumination

A number of different bright-field illumination configurations were used to create the MCAM data presented here. For the freely moving zebrafish and C. elegans results, we relied on a set of 4 different 32×32 LED arrays, each 13 x 13 cm in size with a 4 mm LED pitch (Adafruit Product ID 607) placed approximately 15 cm beneath the sample plane, with a diffusive screen placed approximately 4 cm above the LED array. While these illumination LED arrays offer individual control over the specific color channel (red, green and blue) and brightness of each LED within the array, the experiments presented here typically illuminated all LED color channels at uniform brightness (i.e., uniform white illumination). For epi bright field illumination e.g., for imaging slime mold, ants, and adult Drosophila, we used LED arrays held off to the side of the behavioral arena by a custom-designed 3D printed adapter specifically built for these purposes. The adaptor was mounted to a plastic flexible gooseneck so we could adjust the light to maximize brightness and minimize glare.

### MCAM Fluorescence imaging setup

Wide-field single-channel fluorescence imaging (**Figure 3**) was implemented with two external high-power LEDs with a wavelength of 470 nm (Chanzon, 100W Blue) for excitation. Each excitation source additionally included a short-pass filter (Thorlabs, FES500). We then inserted a custom-designed emission filter array containing 24 filters (525±25.0nm, Chroma, ET525/50m) directly over the MCAM imaging optics array to selectively record excited green fluorescent protein and GCaMP. Dual-channel fluorescence imaging was accomplished with a set of blue LEDs (470 nm, Chanzon, 100W Blue) combined with a 500 nm short-pass filter (Thorlabs, FES500). For emission, we used 510±10.0 nm filters (Chroma, ET510/20m) for *Tg(elavl3:GCaMP6s)* (green) signals and 610±37.5nm filters (Chroma, ET610/75m) for slc17a6b (red) signals. Both channels are simultaneously captured for 240 seconds, then each channel is stitched separately using PTGui software. Fish were segmented from stitched images using OpenCV functions and tracked manually. Fluorescent activity per frame was calculated by averaging intensity within each segmented fish brain area for both green and red channels and computing subsequent ΔF/F ratiometric measurement. The speed of the fish was measured by calculating the distance of the center point of the segmentation mask from each frame.

### MCAM Software

#### Acquisition

All data acquisition was performed using custom-developed Python software. Various Python scripts for controlling image acquisition (e.g., exposure time, gain, number of micro-cameras for image capture, video acquisition parameters) were both available for command-line execution and integrated into a custom PyQT graphical user interface. Python scripts were also developed to route and organize acquired image and video data within memory. Once setup, video or images data can be captured automatically for hours without human supervision. Additional data management details are in Supplementary Materials: MCAM Data Management.

#### Image preprocessing

Before analyzing full-field image data, we first applied a flat-field correction to each micro-camera image to remove vignetting and other artifacts caused by variations in the pixel-to-pixel sensitivity of individual micro-cameras. This correction was performed by normalizing each pixel to the value of the mean of a set of five out-of-focus full-field images of a white diffuser captured at the end of each experiment. After this standard flat-field correction, we then performed image stitching on a per-frame basis as explained below.

#### Image stitching

For finalized, multi-camera images, we either used commercial stitching software PtGui (Rotterdam, The Netherlands), or a custom Fourier stitching algorithm to combine the images into a final stitched panorama. For Fourier stitching, shifts in the spatial domain correspond to a change in slope in phase in Fourier domain which becomes a matched parameter. Hence, stitching parameters were acquired by first computing the Fourier transform of every image from each CMOS sensor, and then calculating phase differences to determine the shift in the spatial domain. As computational stitching algorithms require features in the image to work successfully, and many of the images of biological samples are relatively sparsely populated with features (e.g., **Figure 2a**). Hence, at the beginning of each imaging session, we acquired images of control target images selected to contain variation in the frequency and phase domains (e.g., pictures of natural scenes, text, or other complex scenes) that cover the entire field of view and use these to generate stitching parameters for the images from subsequent experiments. See Supplementary Materials: Image Stitching for additional detail.

#### Depth detection

The MCAM system was configured to have slightly more than 50% FOV overlap in adjacent horizontal cameras to enable the system to image an object with at least two cameras at any given timepoint. Stereoscopic depth tracking was the achieved with custom-written Python software (see details in Supplementary Materials: Depth Tracking). Briefly, following object detection (outlined below), features were identified in stereo-image pairs of objects of interest from neighboring micro-camera pairs with standard OpenCV packages. Feature pairs were identified via the minimum Hamming distance between the per-image lists of candidate feature vectors. The average distance between feature pairs was then used as a measure of object disparity at the sensor plane, from which object depth is established via a geometric relation.

#### Bright field imaging

##### In freely swimming zebrafish larva

In-vivo light field imaging in the MCAM system was performed in wild-type, tüpfel long fin zebrafish (TL) at 6-8 days-post-fertilization. Zebrafish were placed in a custom 3D-printed arena filled with E3 medium to a height of 5 mm. Arenas were designed using Fusion360 (Autodesk; CA, USA). For gigapixel imaging, the arena was rectangular with curved corners, with dimensions 11 cm x 22 cm, wall thickness 1.5 mm, and height 5 mm. The arena was printed with black PLA (Hatchbox, CA, USA) and glued onto a glass sheet using black hot glue around the outside of the arena. For data shown in **Figure 1**, we recorded spontaneous swimming behavior of 130 larvae over 1 hour. All data acquisition and preprocessing was performed using custom Python software.

#### Object detection

To detect fish in stitched images generated from stitched images, we trained a Faster-RCNN object detection network (**Supplementary Figure 5A**) using *TensorFlow* using Google’s object detection API. For more details about the network, training, and parameters used, see discussion of **Supplementary Figure 5** (Ren et al., 2016). Briefly, this deep neural network takes in an arbitrary image and generates bounding boxes (and confidence values) around all putative fish in the image (**Figure 2A**). We made several modifications to standard training and inference stages, including adding a novel occlusion step to the augmentation pipeline (**Supplementary Figure 5D**). Because the images are much too large for training and inference using standard GPU RAM, we broke up the training images into patches that contained the objects of interest (as well as negative examples, such as debris) and used these sub-images for training (**Supplementary Figure 5B**). For inference, we use a moving window followed by non-max suppression to remove redundant detection events (**Supplementary Figure 5H**). A repository where users can find the test the frozen *gigadetector* network, and apply it to new images, can be found at GitHub at https://github.com/EricThomson/gigadetector.

#### Zebrafish Identification

To demonstrate the resolution of MCAM, we used it to capture video of nine zebrafish larvae and trained a convolutional neural network (CNN) with images obtained from the video’s frames to distinguish between detected fish. We devised the CNN in *Tensorflow* as a siamese neural network with triplet loss (Schroff et al., 2015). The architecture consists of three sets of two convolutional layers with rectified linear unit (ReLU) activation functions and 1 max pool, and a final two fully connected layers. The last fully connected layer outputs 64-dimensional embeddings (**Supplementary Figure 6B**). Inputs to the network were a set of 3 images: two images of the same fish taken from different frames (*f*_*i*_ and 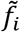), and one image of a different fish (*f*_*j*_). The network was then optimized during training to minimize the Euclidean distance between embeddings of *f*_*i*_ and 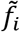 and maximize the Euclidean distance between embeddings of both *f*_*i*_ and *f*_*j*_, and 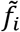 and *f*_*j*_. We performed a temporal validation of the network, with the first 80% of frames used for training and the last 20% for testing. After data augmentation we generated 250 images per larval zebrafish, with the 80/20 split resulting in 1800 training images and 450 testing images. We trained using the Adam optimizer (with default parameters) for 62 epochs with a batch size of 32. To reduce computational complexity, we performed 2x image down sampling to 224 × 224 pixels; at this resolution, feature differences across fish, including variegated melanophore patterns, are still apparent. We used t-SNE (Maaten and Hinton, 2008) to visualize the 64-dimensional embeddings produced as output from the network. For additional details see Supplementary Methods.

##### Caenorhabditis elegans

*C. elegans* strains were kindly provided by Dr. Dong Yan, Duke University, by way of the Caenorhabditis Genetics Center (CGC) at the University of Minnesota. We used three different strains. One, a wild-type (N2) line. Two, a unc-3 mutant CB151 that is a loss of function mutant. Three, a novel transgenic line NYL2629 that is a cross between strains CZ2475 [Pflp-13∷GFP(juIs145) II and OH11746](Donnelly et al., 2013) and OH11746 [Punc-3∷mCherry+pha-1(otIs447) (Kerk et al., 2017) that express fluorescent proteins in two sets of motor neurons. To grow the colonies, we followed standard procedures (Stiernagle, 2006). Briefly, we prepared nematode growth medium (NGM) plates using medium acquired from Carolina Biological (NC, USA). We melted the solid NGM in a hot water bath at 95°C, and before pouring, we cooled the bath to 55°C to minimize warping of the petri dishes and condensation. We used a large petri dish (24.5×24.5cm Square Bio Assay Dishes; Thermofisher). To prevent contamination, we flamed the bottle lip before pouring. Once the NGM was cured (about an hour), we then grew a lawn of *E coli* in the medium by inoculating the NGM with OP50 and letting it grow overnight in an incubator at 37°C. OP50 was kindly provided by Dr. Dong Yan at Duke University. To grow a colony of *C. elegans*, we transplanted chunks of pre-existing lines to the *E. coli* lawns and monitored the dish until we observed adult *C. elegans* moving through the lawn on the majority of the dish (typically 2-4 days).

#### Physarum polycephalum

Desiccated slime mold (*P. polycephalum*: Carolina Biological; NC, USA) was maintained at room temperature until ready for use. To activate the sclerotium, we set it on a plate of agar (1-4%; Becton, Dickinson and Company; NJ, USA), and hydrated it with a drop of distilled water. We liberally sprinkled raw oatmeal flakes on the agar. Under these conditions, the slime mold enters the plasmodium stage, a moist amoeboid state that grows in search of food in its environment. Once the plasmodium has grown, we then used it to seed additional agar plates, either directly with an inoculating glass, or by ‘chunking’ a volume of agar and placing it on a second agar plate, as we did to transfer *C. elegans*.

We used 3D printing to create plastic mazes, which we designed using Fusion360 (Autodesk; CA, USA). The mazes were square (16.5cm outer diameter), with wall thickness 3 mm, height 4mm, and passage width of 12 mm. Mazes were printed with cool gray PLA (Hatchbox, CA, USA). Topographically, the maze was symmetric in that there were four starting positions, one in each corner, and for each starting position the slime mold would have to travel the same minimum distance (25 cm) to the final central position (**Figure 5A**). To construct the maze, we poured 2% agar into a large square (24.5×24.5 cm) petri dish (Nunc Bioassay Dishes; Thermofisher, MA USA), and then placed the maze into the agar while it was still in its liquid phase. We found embedding the maze in the agar better than placing it on top of cured agar, because the slime mold typically grew underneath any items placed on the agar.

After setting up the maze, but before placing the slime mold inside, we disinfected the maze and oatmeal flakes with UV light for three minutes. It is important to keep the petri dish covered during long recordings, so the agar does not dry out. We prevented condensation on the cover by coating it with an anti-fog spray (Rain-X; TX, USA).

Spreading unscented aluminum antiperspirant gel (Dry Idea; Henkel, Düsseldorf, Germany) on top of the maze was an effective deterrent to “cheating” behavior (crawling over the top of the maze). We found this by trial and error with many toppings that included nothing, Chapstick (menthol/strawberry/cherry) (ChapStick; NC, USA), Campho-Phenique (Foundation Consumer Healthcare; PA, USA), and scented antiperspirant (Degree; Unilever, London, U.K.). We observed that the latter two substances were so aversive that the slime mold tended to simply stop growing even within the main passages of the maze.

##### Camponotus pennsylvanicus

Black carpenter ant (*Camponotus pennsylvanicus)* workers were acquired from Carolina Biological, NC. The ants were housed in colonies of ~50 individuals and fed with a mixture of leaves and wood provided by Carolina Biological. The colonies were kept dormant in a refrigerator at ~4 degrees Celsius. Prior to experiments, the colony was removed from the refrigerator and kept at room temperature for at least one hour until ants exhibited baseline levels of movement. To observe the responses of groups of ants to different levels of light, approximately 20 ants were transferred from the colony into a rectangular 3d printed enclosure. The group was imaged for 2 minutes under visible light, which was turned off for 2 minutes. This cycle was repeated 6 times. The entire time, the ants were illuminated from above by IR light sources.

#### Fluorescence imaging

##### Zebrafish

For all experiments, we used zebrafish 5-10 days post-fertilization (dpf). All zebrafish were either wild-type (TL) or transgenic nacre −/− (*mitfa*) zebrafish; these mutants lack pigment in melanocytes on the skin but not in the eye. Zebrafish were maintained on a 14 hrs. light /10 hrs. dark cycle and fertilized eggs were collected and raised at 28.5 °C. Embryos were kept in E3 solution (5 mM NaCl, 0.17 mM KCl, 0.33 mM CaCl2, 0.33 mM MgSO4). All experiments were approved by Duke University School of Medicine’s standing committee of the animal care and use program. Imaging experiments in this study were performed on transgenic zebrafish *Tg(isl1a:GFP)*(Higashijima et al., 2000), *TgBAC(slc17a6b:LOXP-DsRed-LOXP-GFP)*(Koyama et al., 2011), and *Tg(elavl3:GCaMP6s)*(Chen et al., 2013), generous gifts from Misha Ahrens and Ken Poss.

##### Agarose embedded zebrafish

*In-vivo* epi-fluorescence imaging in the MCAM system was performed in transgenic zebrafish *Tg(elavl3:GCaMP6s)* (Chen et al., 2013) 5-7 days post fertilization. Zebrafish were homozygous for GCaMP6s and nacre, *mitfa* (Lister et al., 1999) to prevent formation of dark melanocytes, effectively clearing the skin. Prior to imaging, zebrafish were embedded in low melting point agarose (2% w/v), which prevented movement artifacts. For these brain-wide neural activity mapping experiments, images were acquired at 10 Hz. For data shown in **Figure 3**, we recorded spontaneous brain activity over 150 seconds.

##### Freely swimming zebrafish larva

*In-vivo* epi-fluorescence imaging in the MCAM system with freely swimming larval zebrafish was performed in transgenic zebrafish *Tg(isl1a:GFP)*(Higashijima et al., 2000), *TgBAC(slc17a6b:LOXP-DsRed-LOXP-GFP)*(Koyama et al., 2011), and *Tg(elavl3:GCaMP6s)*(Chen et al., 2013) 5-7 days-post-fertilization. Zebrafish were placed in a custom 3D-printed arena filled with E3 medium to a height of 1-2mm. Arenas were designed using Fusion360 (Autodesk; CA, USA). For fluorescence imaging, the arena was rectangular with curved corners, with dimensions 9cmx7cm, wall thickness 1.5mm, and height 5mm. The arena was printed with black PLA (Hatchbox, CA, USA) and glued onto a glass sheet using black hot glue around the outside of the arena. For data shown in **Figure 3E-I**, 3-14 zebrafish were placed in the arena and spontaneous activity was recorded for up to 10 minutes. Images were collected at 1Hz.

##### Drosophila melanogaster

Flies (genotype: w; UAS-CD4tdGFP/cyo; 221-Gal4) were kindly provided by Dr. Rebecca Vaadia. Flies were raised with standard molasses food at room temperature. For fluorescent imaging, 3rd-instar larvae were transferred from the fly food to a drop of water using wood applicators. A drop of water was used to constrain their moving area (decrease forward movement and increase rolling behavior to visualize the side view). For adult imaging, a white 3D-printed arena (dimensions 12cm x 20cm, wall thickness 1.5 mm, and height 5 mm) was set between two glass plates.

##### Statistical Analysis

All values reported as mean ± sd unless otherwise stated. In bar graphs, error bars correspond to standard deviation unless otherwise stated.

## Data Availability Statement

One-hour long gigapixel dataset of zebrafish of stitched video will be made available from the authors upon publication. Other datasets generated during the current study are available from the corresponding authors on request. Additional videos of gigapixel recordings are viewable at https://gigazoom.rc.duke.edu/

## Code Availability Statement

All code used for analysis was made using Python or standard approaches in MATLAB 2017b and open-source code extensions and are available from the corresponding authors upon reasonable request. The MCAM object detection code is available at https://github.com/EricThomson/gigadetector. Due to the enormous size of the imaging data sets only example data will be made available online and full data sets by request. Instructions for running the code and reproducing published results is available via the code repository README (rendered by GitHub as a web page) and via Jupyter Notebooks, which include code to produce figures.

## Acknowledgements

We would like to thank the Duke School of Medicine as well as Ramona Optics Inc. for technical support, as well as Maxim Nikitchenko for helpful comments and discussions. Thanks to the Ken Poss lab for the *Tg(islet1:GFP)* fish line, Misha Ahrens for the Tg(elavl3:GCaMP6s) and *TgBAC(slc17a6b:LOXP-DsRed-LOXP-GFP)* lines. We further thank R. Yang for advice and gifts on Drosophila and Wesley Grueber and Rebecca Vaadia for advice and gifts of the transgenic Drosophila lines and Rebecca Yang for assistance with their husbandry, Jason Liu for help with experiment design and 3D printing, as well as Dr. Benjamin Judkewitz and Dr. Mike Orger for early project assistance. We furthermore thank Dong Yan, Albert Zhang, and the Center for Caenorhabditis Genetics Center (CGC) for the C. elegans lines and help with setting up C elegans. J. Burris, S. X and K. Olivera for zebrafish husbandry. This work is supported by grants from N.I.H. (BRAIN initiative R34-EB026951-01A1, SBIR R44-OD24879-02, SBIR 1R44CA250877-01, SBIR 1R43EB030979-01), NSF Award 2036439, Alfred P. Sloan Foundation, and by the Whitehall Foundation. We thank Kristian Herrera for 3D renderings in the demo movie.

## Author contributions

E.E.T., E.A.N. and R.W.H. conceived of this project. R.W.H and M.H. designed and built the imaging system with assistance by J.P. R.W.H., M.H. and J.P. performed experiments to validate resolution, image data acquisition rates, and stitching performance. K.K. and P.C. developed fluorescence imaging hardware. E.E.T., R.B., C.S., Y.C., and K.K. performed the biological experiments. E.E.T., E.A.N, W.J., S.X, C.C. and K.K. wrote analysis software to locate and analyze organisms and to analyze neural activity with help from T.W.D. All contributed to hardware control and data processing, and M.H. wrote the stitching algorithms. S.S. and P.C. contributed to figure preparation. E.T, R.W.H. and E.A.N. wrote the manuscript with input from all authors. E.A.N. and R.W.H conceived and led the project.

