## Supplementary Methods and Materials for "Gigapixel behavioral and neural activity imaging with a novel multi-camera array microscope"

### Supplementary Materials

***MCAM hardware***

**MCAM-24:** As described in the main text, the multi-camera array microscope (**Supplementary Figure 1A**) consists of an array of cameras, each composed of a lens, sensor, and optomechanical housing. Each camera images a unique area of a large sample plane in parallel with the other cameras. The base unit of an MCAM array consists of 24 cameras arranged in a 6x4 array (an MCAM-24: **Supplementary Figure 1A**). The 24 CMOS image sensors (1.4 µm pixel, 10 Megapixel, Omnivision OV10823) that comprise an MCAM-24 are integrated onto a single PCB which is controlled by custom-developed electronics (4 follower FPGAs and 1 leader FPGA) to route data via USB 3.0 to a desktop computer. The sensors are arranged at a 19 mm pitch. Customized mounts fit over the sensor array and attach to a 24-lens adjustable mount, which holds a 6x4 array of individually focusable M12 lenses (f=25 mm, Edmund 58-207) at 19 mm pitch which are focused at a 150 mm object distance (**Supplementary Figure 1B**). The resulting raw image data from this arrangement (**Supplementary Figure 1C**) contains 240 Megapixels (MP), which is then sent to our image stitching and analysis software to create the final resulting images and video (**Supplementary Figure 1D**).

**MCAM-96:** To create the gigapixel multi-camera array system in **Supplementary Figure 1E**, we combine 4 of the MCAM-24 systems into a 2x2 configuration, via the use of optomechanical mounting plates (**Supplementary Figure 1F**). This results in an 8x12 array of lenses and sensors, with 4 sets of electronics to send image data via 4 USB 3.0 lines to a single desktop computer. The MCAM-24 arrays were designed to ensure that 19 mm inter-sensor spacing is maintained when 4 individual systems are joined together into an MCAM-96 arrangement. Similar image stitching and analysis software is then applied to each of the 960 MP raw captured frames to create the final video (**Supplementary Figure 1G**).

**Illumination:** LED arrays were used to illuminate samples either from above (epi-illumination), from below (trans-illumination), or both. When using trans-illumination, one and four commercially available LED arrays (32x32 RGB LED Matrix, Adafruit) were used for the MCAM-24 and MCAM-96 setup, respectively. In some experiments, we placed a sheet of diffusive paper over the LED array to provide more even illumination. For epi-illumination, a custom-developed LED array was used that contained approximately 500 LEDs arranged on a PCB around a 4x6 grid of 19 mm holes within the board, for each micro-camera lens to image through.

**Supplementary Figure 1**

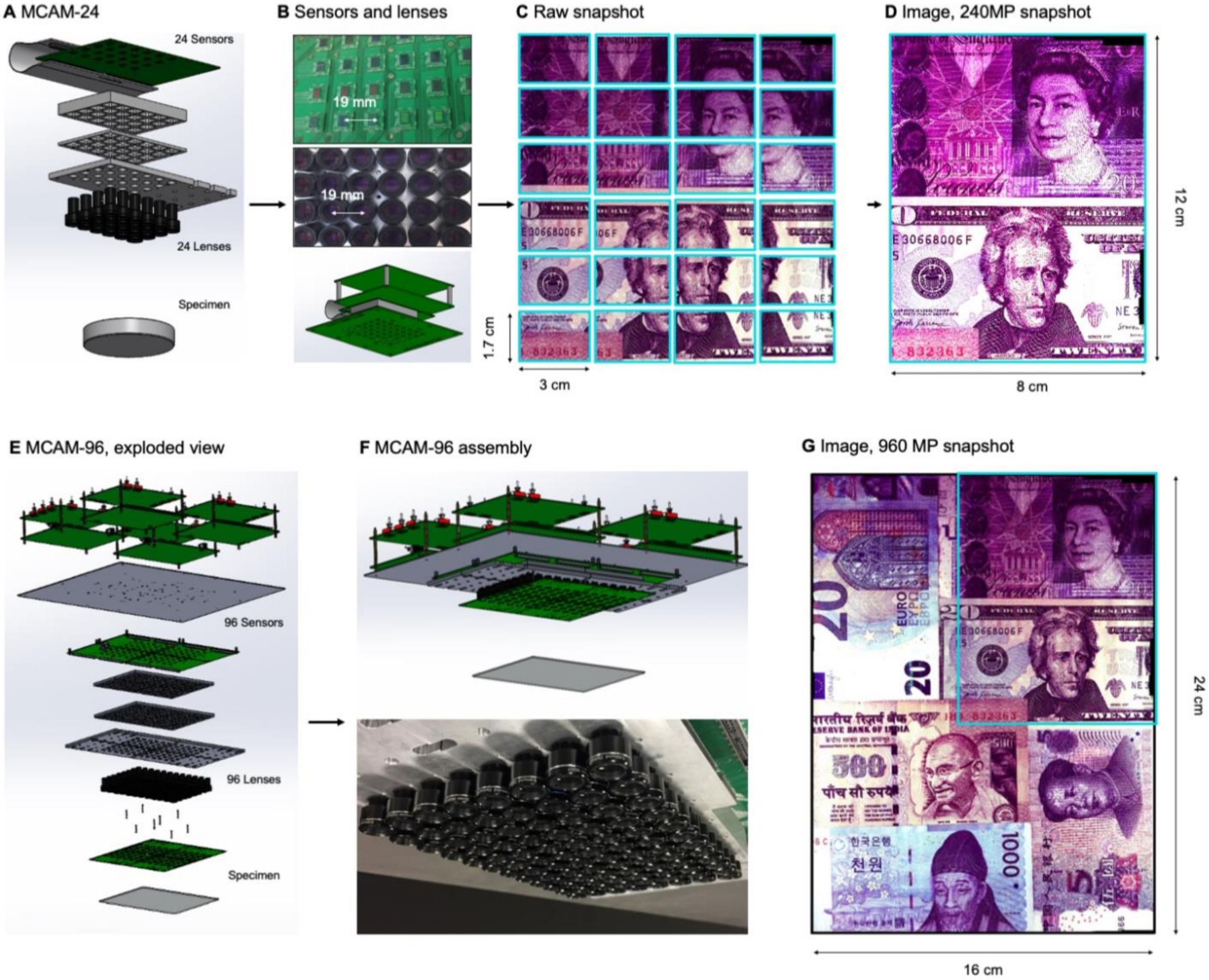

**Supplementary Figure 1 | MCAM hardware.**

**A.** MCAM-24 hardware arrangement in an exploded view, showing an array of 24 individual CMOS sensors (10 MP each) integrated onto a single circuit board with 24 lenses mounted in front.

**B.** Photos of bare MCAM-24 CMOS sensor array and lens array.

**C.** Example snapshot of 24 images acquired by MCAM-24 of common currency.

**D.** Stitched composite.

**E.** Four MCAM-24 units are combined to create the gigapixel MCAM-96 setup, with 96 sensors and lenses tiled into a uniformly spaced array for a total of 960 MP captured per image snapshot.

**F.** CAD render and photo of complete MCAM-96 system.

**G.** Example stitched composite from MCAM-96, with MCAM-24 field-of-view shown in teal box.

**MCAM resolution analysis and verification**

The MCAM imaging system employed optics that were designed to provide an object-side numerical aperture (NA) of approximately 0.03 (9 mm optical lens diameter at an approximately 150 mm working distance). Under incoherent illumination (with wavelength λ = 500 nm), this NA leads to an incoherent transfer function cutoff spatial frequency of $f_c = 2NA/\lambda = 0.12\ \mu m^{-1}$ at the object plane (120 line pairs per mm). This corresponds to a two-point Sparrow resolution limit cutoff of $d_{min} = 0.47\ \lambda/NA = 7.8\ \mu m$ under incoherent illumination, and $d_{min} = 0.73\ \lambda/NA = 12.2\ \mu m$ under coherent illumination (Ou et al., 2015) at the object (i.e., sample) plane. The use of relatively small micro-camera lenses in our experimental system led to minimal aberrations across each camera field-of-view, and thus minimal aberrations across the entire MCAM field-of-view.

The lens focal length (f=25 mm) was selected to produce an imaging system magnification $M = d_i/d_o = 30\ mm / 150\ mm = 0.2$ to fulfill our tiled field-of-view specifications. With an imaging sensor pixel size of 1.4 µm, we were able to sample at nearly this maximum achievable optical resolution limit. In other words, the imaging system resolution (optical and pixel spacing) was designed to be at the limit of critical sampling for coherent imaging. With a 0.2 imaging system magnification, the pixel size projected into the sample plane is 7 µm for the MCAM, which is close to half the coherent imaging Sparrow limit noted above.

Our final demonstrated resolution was close to both the expected diffraction-limited and pixel-limited system resolution. As sketched in **Supplementary Figure 2A-B**, we used the lens, sensor and spacing specifications outlined above to create an arrayed system to capture image data from across a continuous field-of-view for subsequent post-processing. By using rectangular image sensors, we were able to create slightly more than 50% overlap along one dimension for depth tracking, as revealed by the MCAM's cross-sectional field-of-view geometry and discussed further in the main text.

**Supplementary Figures 2C-D** show an MCAM full-field image of a custom-designed resolution target (Photo Sciences Inc, https://www.photo-sciences.com/) spanning its full 18 x 24 cm imaging area. Zoom-ins highlight small, custom-printed US Air Force targets that allow us to assess the impact of aberrations from randomly selected areas. Using this custom-designed target, we were able to experimentally verify a resolution cut-off that consistently lies between group 5 element 5 and 6, which exhibit full-pitch resolutions of 17.54 µm and 19.68 µm, respectively, leading to our claim of 18 µm resolution in the main text. This experimental resolution is slightly worse than the theoretically predicted full-pitch resolution and can be attributed to mild aberrations and the potential contribution of slight defocus that can vary as a function of field-of-view position. Finally, **Supplementary Figures 2F** shows images of 5 µm and 3 µm polystyrene microspheres captured with both our MCAM and a standard 20x microscope as comparison. We observe that the MCAM can reliably detect the 5um spheres, and consistently detects the 3 µm microspheres but not at full fidelity. Accordingly, we have estimated MCAM sensitivity at 5 µm.

Future MCAM designs can easily achieve higher imaging resolution. A tighter inter-camera spacing can lead to a larger per-camera image magnification, which in turn can increase their maximum spatial resolution. We selected an inter-camera spacing of 19 mm here and used imaging lenses that had an 18 mm maximum outer diameter. The use of alternative lenses can lead to tighter packing. A smaller sensor pixel size will be used in future system designs, along with a tighter micro-camera spacing, to achieve a higher imaging NA and resolution. In addition, we did not account for the effects of a Bayer filter over the sensors' CMOS pixels in the above analysis. We used image normalization techniques to remove the effects of the Bayer filter for our grayscale resolution tests. The inclusion of a Bayer filter over each CMOS pixel array further limits detector resolution when utilized to produce snapshot color images. Furthermore, the Bayer filters also reduce sensor sensitivity in general. For applications that could benefit from higher spatial resolution, future designs will use unfiltered monochrome pixels.

**Supplementary Figure 2**

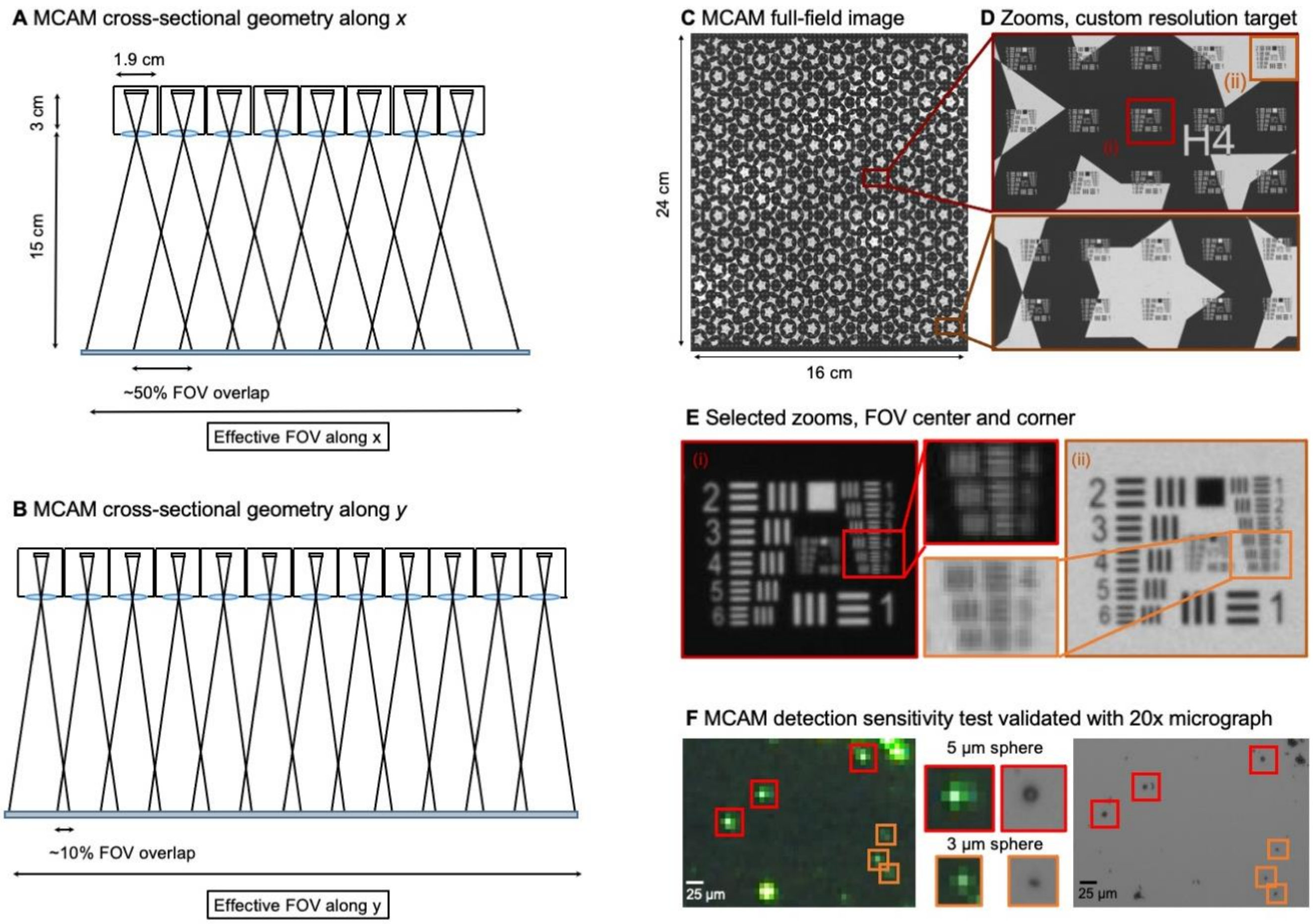


__**Supplementary Figure 2 | 96-camera MCAM geometry, resolution measurements and sensitivity test.**__

**A.** MCAM geometry cross-section along x direction of 8 x 12 array. The rectangular image sensor exhibits a longer dimension along x, yielding camera images sharing >50% FOV overlap for depth tracking and dual-fluorescence imaging.

**B.** MCAM geometry cross-section along y direction of 8 x 12 array, where image sensor's shorter dimension leads to approximately 10% overlap between adjacent camera FOVs to facilitate seamless image stitching.

**C.** Full field-of-view MCAM-96 image of a custom-designed resolution target covering over 16 cm x 24 cm, with **D.** Zoom-ins.

**E.** Two further zoom-ins from marked boxes in (d) that lie at the center and edge of a single camera FOV. Resolution performance varies minimally across each camera FOV. For this custom-designed resolution target, Element 4 at the resolution limit exhibits a 22.10 µm full-pitch line pair spacing, Element 5 a 19.68 µm full-pitch line pair spacing, Element 6 a 17.54 µm full-pitch line pair spacing. The MCAM consistently resolves Element 5 but at times does not fully resolve Element 6, leading to our approximation of 18 µm resolution.

**F.** MCAM sensitivity imaging experiment. The left images show 3 µm and 5 µm diameter microspheres (Polysciences Polybead) deposited on mirrored surface and illuminated with a mounted LED (505 nm, Thorlabs M505L4) at 45-degree angle for dark-field imaging. Red boxes mark location of 5 µm spheres, orange arrows mark 3 µm spheres. On the right are the same region of the specimen imaged with a 20X microscope, verifying presence of 3 µm and 5 µm single microspheres. The MCAM reliably detects 5 µm single microspheres and, as shown, can detect 3 µm microspheres, but at times does not capture sufficient signal from the 3 µm features.

***Image stitching***

The MCAM relies on image stitching software to combine images from all micro-cameras into a final integrated composite image (a 'stitched' image). Stitching is not required for many applications (e.g., organism detection and cropping, organism tracking, fluorescence analysis), as the application-specific software can be applied directly to the raw image data. However, for applications such as image segmentation, or to produce finalized, multi-camera images for visualization or presentations, stitching becomes an important and helpful post-processing step.

The general workflow of our stitching process is outlined in **Supplementary Figure 3A**. In this work, we relied on two unique stitching algorithms written in Python but applied the same workflow to each. In short, we took advantage of calibration targets to help establish accurate stitching parameters for our experimental data, which we then applied to produce our final stitched video frames.

1. **Calibration target**: Immediately following the image or video data collection step for our different sets of experiments, and before altering the position of the camera lenses or sample plane (i.e., before moving anything except the sample), we captured 2-4 images of a calibration target. The purpose of calibration target imaging is to acquire stitching parameters that may be otherwise challenging to extract from the raw experimental data, because the experimental data often did not contain a rich set of features (a few fish against a blank background). As detailed below, image stitching algorithms perform best with targets that include a large range of non-zero spatial frequencies across the sample plane. For instance, pieces of paper covered with text and wavy lines, photocopies of full-page photographs, or our custom-designed resolution target all worked well as calibration targets.

2. **Parameter extraction**: Before stitching each dataset, we processed the MCAM image set of the calibration target acquired in step 1 to extract key parameters. Details of the processing steps using each stitching approach are detailed below. Each approach generates a stitched calibration target image, as well as a set of parameters (e.g., a set of coordinates (x,y) to place each image, and/or any additional affine transform parameters such as per-image shearing and rotation), which we saved for later use. We visually assessed the stitched calibration target images and, if any issues were identified (rarely encountered in later experiments), we would alter stitching algorithm hyper-parameters (discussed below) and re-run.

3. **Final stitch**: After extracting a set of parameters for the MCAM image set (24 or 96) of the calibration target, we used the same software to apply the saved stitching parameters to stitch the experimentally acquired MCAM image data, see example in **Supplementary Figure 3B**. Video data was stitched on a per-frame basis.

We use two different stitching methods at different stages of our experimental development and testing.
**Fourier stitching algorithm:** Fourier-domain image stitching is a well-established method to efficiently align different overlapping image segments (Szeliski et al., 2006). Briefly, shifts in the spatial domain correspond to a slope change of the phase component of an image's Fourier domain representation (i.e., its 2D spatial frequencies). Accordingly, by digitally computing the 2D fast Fourier transform of each image and determining the phase slope variation between them by computing their normalized product (i.e., phase correlation (Szeliski et al., 2006), one can then extract relative spatial image shifts by performing an inverse Fourier transforming the result. Once relative shifts are identified, images can be appropriately aligned and an error metric applied to examine how well the overlapped image areas match, which can subsequently be minimized as a function of the shift variables. This type of "direct" stitching approach compares images via pixel-to-pixel matching, instead of first searching for specific features, and provides a robust and relatively rapid approach.

**PtGui:** We also adopted a commercial stitching software platform, PtGui (Rotterdam, Netherlands), to stitch MCAM images. This software comes with a simple batch processing mode, so that once you extract the stitching parameters for one calibration target, you can then apply the same parameters to all of the images in a directory, which we found it to be extremely flexible, accurate, and fast.

**Supplementary Figure 3**

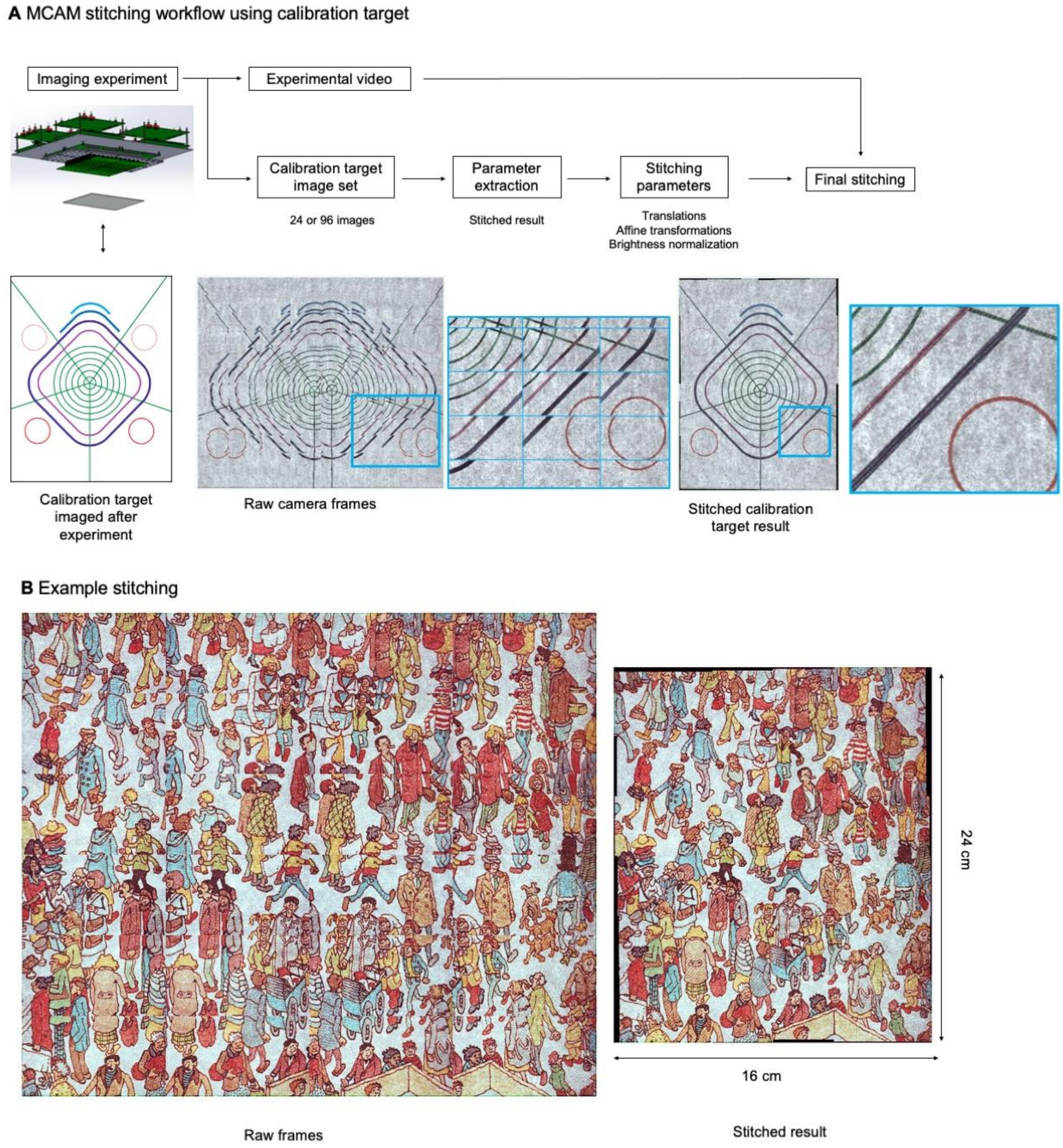

**Supplementary Figure 3 | MCAM Stitching Process.**

**A.** Schematic of workflow of the MCAM stitching process.

**B.** Example raw frame are stitched together using parameters extracted from calibration target.

##### *MCAM Data Management*

The outline of MCAM image data management is diagrammed in **Supplementary Figure 4A**. MCAM-96 comprises four MCAM-24s, each of whose sensors are integrated on a common PCB board. Data from each set of 24 cameras is routed to a single USB 3.0 cable via a leader-follower FPGA configuration, yielding 4 USB 3.0 lines that connect the gigapixel system to a single desktop computer. During video capture, approximately 240 MB/sec of image data is sent along each USB 3.0 line for a 1 GB/sec approximate total data rate, which is subsequently saved in solid-state memory.

Due to limited data transmission speeds, our "full-frame" image capture and transmission strategy, delivering data from all of the cameras, utilized frame interleaving from consecutive cameras within each 24-unit array. The resulting timing sequence of this interleaved arrangement is sketched in **Supplementary Figure 4B**. Every 1 second (which defines the overall system frame rate), one image from each of the 24 cameras is exposed and then routed via the FPGAs to computer memory in a staggered manner. While the per-sensor exposure time can exceed 42 ms (i.e., 1/24 of a second), the delay between captured snapshots from one camera to the next is set (approximately) at this quantity. This type of timing sequence is followed by each of the four 24-unit MCAMs comprising the full gigapixel system.

In an alternative configuration, the MCAM system can also be operated in a "single-frame" image capture mode, where image data from any one of the 24 cameras can be streamed at approximately 10-15 frames per second, and the user can programmatically switch which camera they are streaming data from with a minor delay (100-200 ms, depending upon exposure and frame rate), as sketched in **Supplementary Figure 4C**. Within the gigapixel MCAM, it is possible to simultaneously stream from up to 4 cameras at this higher frame rate. By providing a 10X higher imaging speed, this second streaming option is helpful for monitoring faster organism dynamics and with system setup and alignment. In the future, it can enable automated tracking and recording of moving objects, where sensors are configured to switch single-frame data streaming on/off as a moving organism, for example, passes within each field-of-view. The resulting dataset will yield a seamless video of the single organism at this higher frame rate, much like a mechanical tracking system (Krishnamurthy et al. 2020), but without requiring any moving parts.

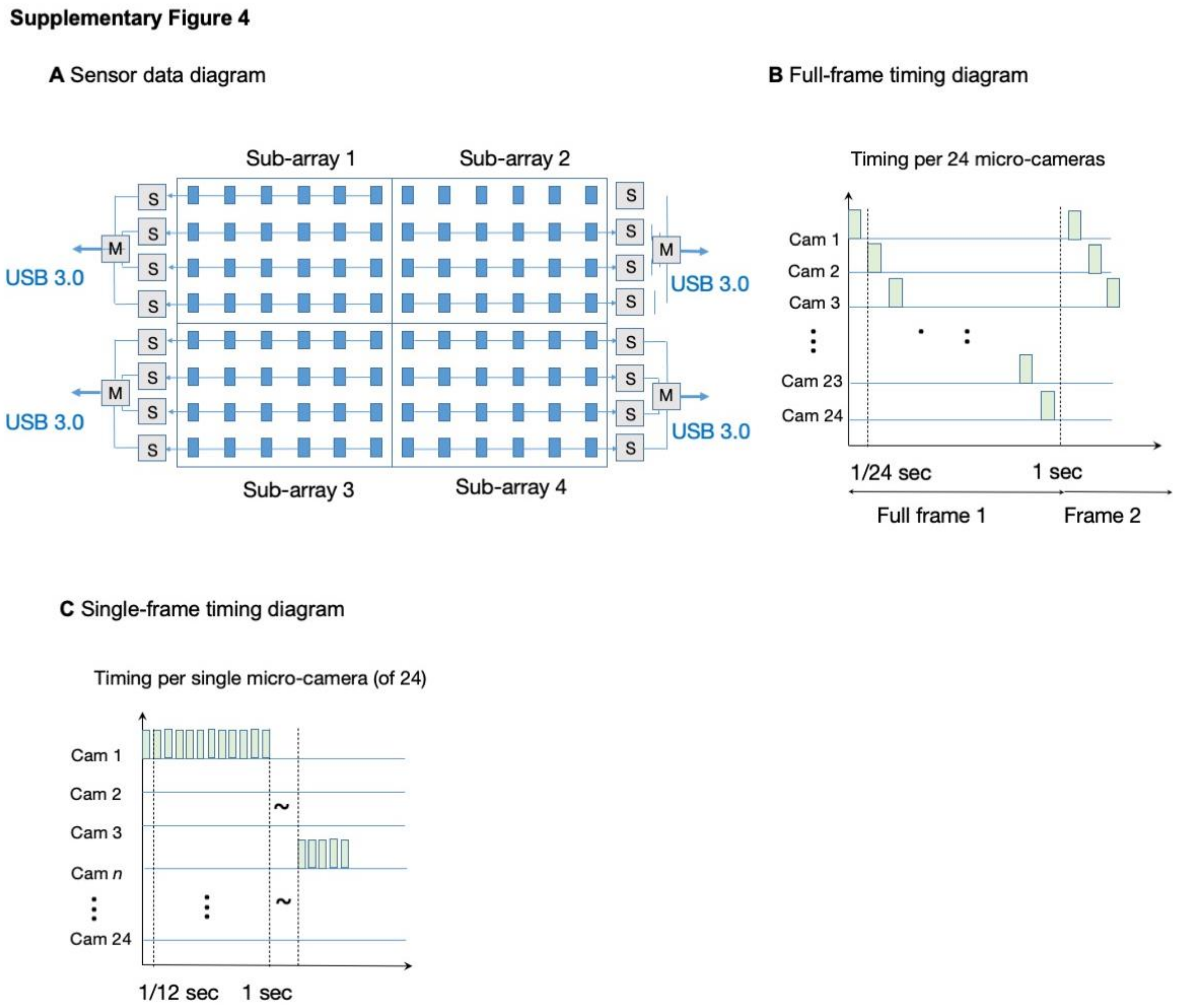


**Supplementary Figure 4 | MCAM data management and timing.**

**A.** MCAM-96 (0.96 gigapixel) data layout and readout geometry from four MCAM-24 sub-arrays.

**B.** Timing sequence of frame data for each MCAM-24 running in full-frame mode.

**C.** Timing sequence of frame data for each MCAM-24 running in single-frame mode.

***Large-scale object detection pipeline***

We adapted a well-known deep-learning based object detection algorithm (Faster-RCNN) to work at gigapixel scale. We used this architecture because of its high accuracy (Huang et al., 2017). The Faster-RCNN architecture has two main components: a traditional Convolutional Neural Network (CNN) that extracts features from the original image, and a region proposal network (RPN) that is trained to find regions in the image that are likely to contain objects Once the RPN suggests good candidate regions to look for objects, then a traditional classifier can be applied to features in those regions (**Supplementary Figure 5A**). We used a Faster-RCNN with a resnet-101 backbone CNN that was pre-trained on the COCO dataset for feature extraction (Lin et al. 2014), and then subsequently finetuned using our data, as described below. In what follows, we provide details on the four main steps required to train and run the network with novel bright-field imaging data. While we discuss our pipeline in the context of zebrafish data, the framework is applicable to any type of organism.

1. Annotation of images
Training a network for object detection requires a set of ground-truth images that contains objects with bounding boxes circumscribing their perimeter. To generate such annotated data, we selected a random sample of frames from our library of movies of freely moving zebrafish. Before annotation, we preprocessed the images by normalizing and stitching together images from individual cameras as discussed previously (**Figure 1** and **Supplementary Figure 3**).

After stitching, the images have dimensions as high as 15000 × 23000 pixels (width x height), which is far too large to fit in GPU memory during training. To reduce peak memory usage for training and inference, we broke up the images into smaller sub-images. Specifically, we selected all 1024 × 1024 sub-images that contained at least one fish as well as *negative* examples such as debris, arena walls, etc. (**Supplementary Figure 5B**).

We annotated these sub-images using the open-source *labelImg* software (https://github.com/tzutalin/labelImg). Annotation consists of drawing bounding boxes around, and supplying the category for, every object of interest. Because each sub-image is relatively small, this process went much faster than the equivalent process of drawing bounding boxes in full-sized MCAM images: an experienced annotator typically annotates more than 100 sub-images an hour. For the negative examples no bounding boxes need to be drawn: the images simply need to be verified as valid.

The goal with annotation was to maximize precision (i.e., minimize false positives), and draw boxes as we hoped the algorithm would learn to draw them. We found that the decisions we made were directly reflected in the performance of the network. Over time, we adopted the following convention: for purposes of annotation, we defined a "fish" as any image of a fish that included the head (which by definition had to include at least 1/3 of an eye down to 3/2 of the distance from the tip of the nose to the caudal tip of the swim bladder). Headless fish tail pieces ("sushi") were not annotated.

We needed to adopt such unusual conventions because of stitching artifacts, which sometimes made annotation challenging. Approximately 7% of training images that contained fish (65/940) included stitching artifacts. These artifacts included duplication of object parts (12/65), translocation of fish (26/65), and complete detachment of segments of zebrafish, so they simply appeared as isolated fish parts (27/65) (**Supplementary Figure 5C**). For translocation artifacts, if the tail piece was detached from the head piece by fewer than 20 pixels, then we circumscribed all the parts as the same fish. If a fish part was duplicated due to asynchronous image acquisition between cameras, we annotated all those individual parts that satisfied the above definition of a fish.

Following the above conventions, we ended up with 1138 bounding boxes containing fish in 1097 sub-images. Of those sub-images, 157 were pure negative examples and 940 contained fish (763 sub-images contained one fish, 157 contained two, 19 contained three, and one sub-image contained four fish).

2. Augmentation of training data

We split the 1097 sub-images into training and testing data (an 80/20 split of 877 training images and 220 testing images). We did not train the network directly on the training images but used an image augmentation pipeline to generate five augmented images per training image for actual training (a total of 4385 augmented images).

We augmented the training data using the open-source Python package *imgaug* (https://github.com/aleju/imgaug). This package lets you arbitrarily transform each image in the training set, while also performing appropriate corresponding transforms of the bounding boxes so that your annotations will remain intact (for instance if you scale the original image, the bounding box will be scaled appropriately). The annotation pipeline consisted of the following operations applied in the following order (with probability that it will be applied given in parentheses): left/right flip (0.5), up/down flip (0.5), rotate by +/-90 degrees (0.5), change brightness between [-30,80] (1.0), scale from [0.75, 1.5] (0.9), sharpen, blur, or neither (1/3 each), motion blur (0.15), rotate (from +/-5 degrees) (0.9), and a custom-built occlusion step (1/3). For examples, see Supplemental Figure 6D. When a value, such as brightness, is given as a range, it was randomly selected from that range with a uniform probability.

Occlusion events were relatively rare in our actual data (7% of bounding boxes overlapped with other boxes), and the overlap was typically small, with a mean Intersection over Union (IoU) between bounding boxes of 10.9 (**Supplementary Figure 5E**). Hence to ensure the network was able to detect fish in those rare cases when there was overlap, we built a custom occlusion augmentation step. Briefly, we artificially generated occluder fish at a random location within the bounding box of the most central fish in the original sub-image. More specifically, we segmented ten fish images by hand in Photoshop, and saved each of them at eight 45-degree angles in an occluder library. The occluder fish image was augmented before being placed on the background, but with its brightness (i.e., mean pixel value in the center of the fish) set to match that of the background fish to give a more realistic appearance. This custom occlusion pipeline was written in Python using OpenCV. You can see two examples of occluder fish added in **Supplementary Figure 5D** (the second and third augmented figures contain occluder fish whose bounding boxes overlap with the original).

3. Training the network

Once we created the set of test images and (augmented) training data, we then encoded the data for training using the Tensorflow Object Detection API (https://github.com/tensorflow/models/tree/master/research/object_detection). Our first step was to encode the annotated data into Tensorflow record files, which contain the image and bounding box data in a byte-encoded format that can be used by the object detection API to speed up processing.

Once the files are encoded, they can be fed to Tensorflow's object detection API to train the Faster-RNN mentioned above. We trained our current Faster-RCNN 75k steps where we set the learning rate to decay during training: it was 0.003 for the first 10k steps, 0.0003 up to step 15k, and then dropped to 0.00003 after step 25k. The network converged relatively quickly (typically within 30k steps: **Supplementary Figure 5F**) but we found that letting it train longer gave better performance on rare events such occlusions, and also the continued decay in the loss function in the test data indicated that we were not simply overfitting the network (**Supplementary Figure 5F**).

4. Inference with the frozen network

Once the RCNN network had undergone transfer learning, the network was frozen and used for inference on new data. Even for this step, the size of the stitched MCAM images was too large for running inference in GPU RAM. Hence, we sliced the large images into 1024x1024 sub-images using a sliding window beginning at the top left corner of the image (with a stride of 512 pixels) (**Supplementary Figure 5G**). We found it important to ensure that each image patch was exactly 1024x1024. So, for instance when the sliding window would have surpassed the right or bottom edge of the image, we shifted the coordinates back to select a full 1024x1024 sub-image rather than a cropped edge. Without this adjustment, we found the recall of the network suffered considerably.

We applied the frozen network to each sub-image and retained those bounding boxes to which the network assigned a confidence above 0.95. This procedure typically yielded multiple highly overlapping bounding boxes around the same organism from multiple sub-images (false positives were effectively non-existent, as we trained the network to be very selective, as discussed above). To filter out repeats, we first applied non-max suppression (NMS) to select the highest confidence bounding box from an overlapping cluster (NMS threshold 0.5, confidence threshold 0.95). However, even after this step, there were sometimes small boxes (below the NMS threshold) that remained as a subset within the larger boxes, so we applied a final subset filter that removed boxes that were contained within other boxes. These steps yielded the final estimate of the object detection pipeline: a set of bounding boxes, with their associated confidence estimates, like those seen in the original paper (**Figure 2A, Supplementary Figure 5H**).

The entire inference pipeline, including the frozen network and some example images and scripts to help people get started using the network, are available online at the gigadetector repository at GitHub: https://github.com/EricThomson/gigadetector.

**Supplementary Figure 5**

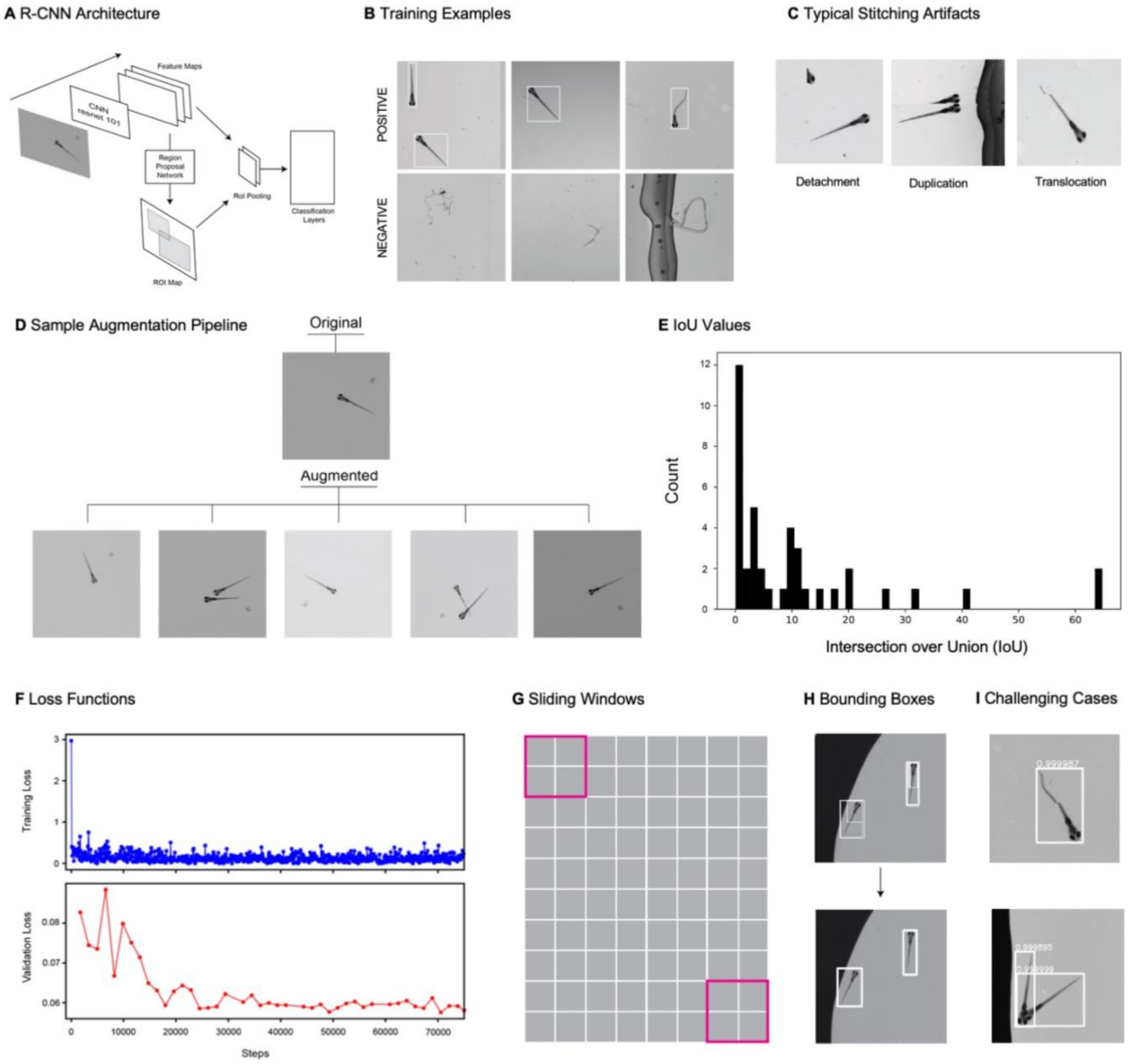

**Supplementary Figure 5| Object tracking using gigapixel video.**

**A.** Faster R-CNN architecture. The architecture consists of a traditional CNN (in this case a resnet-101) for feature extraction, which feeds into a region proposal network (RPN) and an RoI pooling layer. The RPN generates an ROI map, proposing locations of objects. The features and object locations are fed to an RoI pooling layer which normalizes the size of each bounding box before the information is fed into a classifier that predicts the object type for each ROI.

**B.** Examples of positive (top row) and negative (bottom row) examples used for training the faster r-cnn.

**C.** Examples of the main types of stitching artifacts, including detachment (left), duplication (middle), and translocation (right).

**D.** Augmentation pipeline example: top image is the original image, and the bottom five images are five augmented instances generated from the original. We generated five augmented instances of each training image in our training data.

**E.** Histogram of intersection over union values for all fish from our actual data (training and test data).

**F.** Loss function in training and validation data during 75000 step training of the faster-rcnn.

**G.** Depiction of sliding windows used for inference over stitched images.

**H.** Example from a fraction of a stitched frame showing multiple fish of the original set of multiple overlapping bounding boxes, and the final set of unique bounding boxes enveloping single fish (obtained using a combination of a nonmax suppression followed by a proper subset filter).

**I.** Examples of successfully detected and boxed larvae in the presence of stitching artifacts (top) and occlusion (bottom). Please see additional examples in Supplementary Video XX and full gigapixel video data with tracking online.

***Convolutional Neural Networks for Fish Identification***

We used a Convolutional Neural Network (CNN) to differentiate among larval zebrafish. Prior deep learning algorithms have distinguished zebrafish at the juvenile and adult stage (Romero-Ferrero et al., 2019), yet these algorithms have not been shown effective on larval zebrafish, which do not differ significantly in size, and canonical striped skin coloration only emerges in the juvenile stage (Singh et al., 2014). However, wildtype zebrafish larvae do exhibit multiple unique differences in features such as brightness, size, and arrangements of their melanophores, the dark-pigmented cells spread across the body (**Supplementary Figure 6, Figure 2B-C**). We aimed to determine whether we could use deep learning to utilize these differences to distinguish individual larval zebrafish, in images obtained from the MCAM.

We trained our Faster-RCNN object detection algorithm on MCAM images of an arena containing nine zebrafish and used the resulting bounding box coordinates to crop out each individual animal. We then augmented the cropped images via rotations of random angle ($\theta \leq 360°$) (**Supplementary Figure 6A**) and generated 250 images per zebrafish or 2250 images total. We used an 80/20 split for training and testing data to obtain 1800 training images and 450 testing images. The 80/20 data split was performed temporally so that training and testing frames came from non-overlapping periods. We halved the resolution of images to improve training speed and found the melanophore patterns were still discernable.

The architecture of our CNN consists of six convolutional layers and two fully connected layers (**Supplementary Figure 6B**). We designed the CNN as a siamese neural network with a triplet loss, based on successful deep learning face recognition algorithms (Taigman et al., 2014, Schroff et al., 2015). The network receives multiple images and learns their similarities, as reflected by the Euclidean distance between images in a lower-dimensional embedding space within the network. Two images of the same zebrafish (anchor and positive) and one image of a different zebrafish (negative) are fed into three subnetworks with identical parameters and weights. Over training, the distance between the anchor and positive input decreases while the distance between the anchor and negative input increases, resulting in clusters containing images of the same fish.

We used a batch size of 32 and Adam optimization (Kingma et al., 2015). We generated 64-dimensional embeddings of the images and used t-distributed stochastic neighbor embedding (t-SNE) to visualize the embeddings in two-dimensions (Maaten et al., 2008). We found that by 62 epochs the embeddings separated in two-dimensional space into nine clusters, each solely containing images of the same fish (**Supplementary Figure 6C, Figure 2B-C**). This demonstrates our CNN was effective at distinguishing among the nine larval zebrafish, presumably due to the MCAM resolving differences in their melanophore patterns (**Figure 2B**) or other anatomical details.

**Supplementary Figure 6**

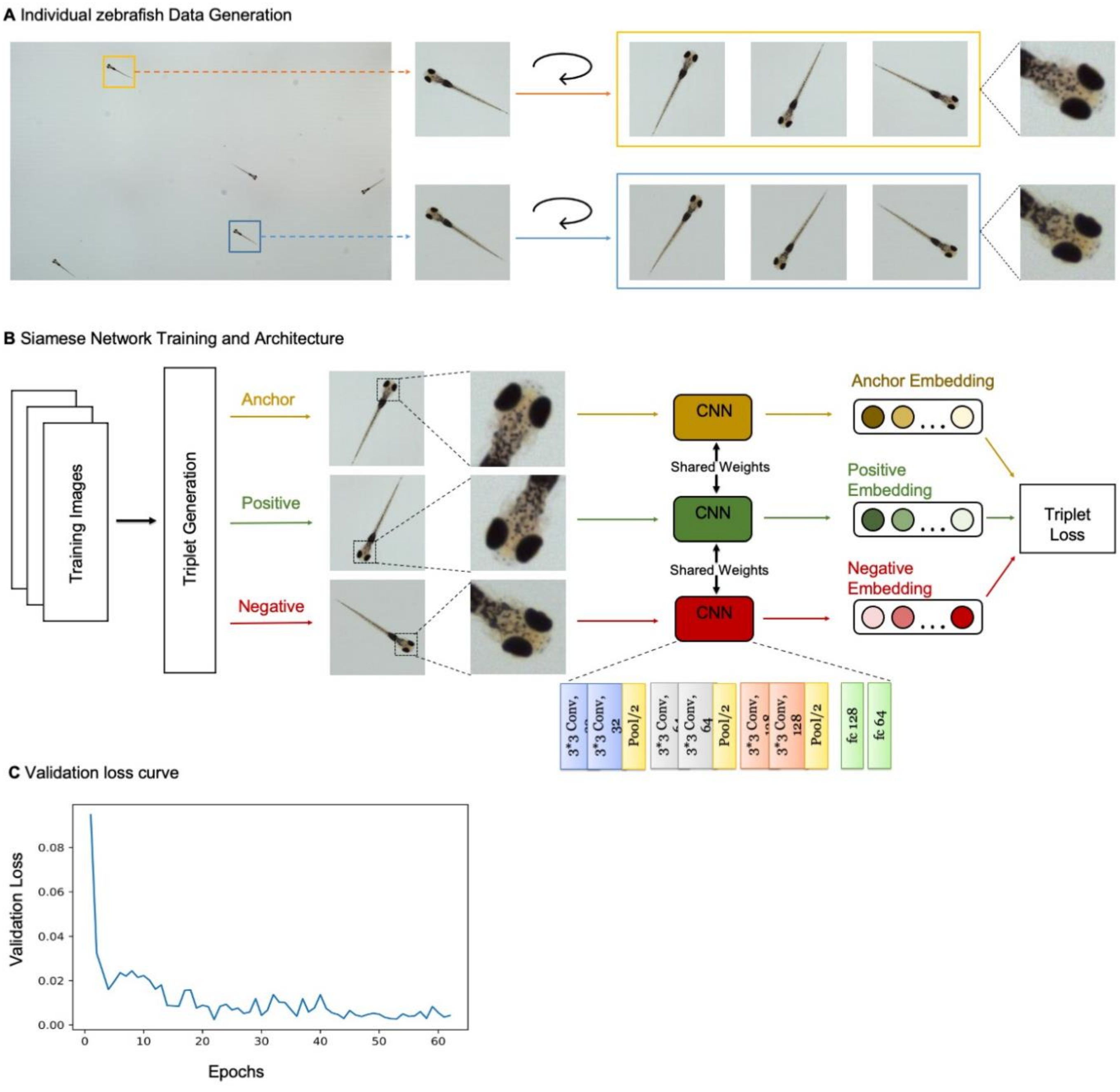

**Supplementary Figure 6| Siamese network with triplet loss differentiates individual zebrafish.**

**A.** Strategy for training data generation from two example larval zebrafish in an arena section. Bounding boxes from Faster-RCNN algorithm are used to crop fish from images captured by MCAM. Data is augmented via rotations of cropped images by a random angle ($\theta \leq 360°$). Melanophore patterns are discernable in training images and presumably are used to distinguish larval zebrafish.

**B.** Cartoon of Siamese Network with triplet loss architecture used to differentiate larval zebrafish using MCAM resolution. Three subnetworks with shared weights and parameters have six convolutional layers and two fully connected layers. The subnetworks receive triplets containing two training images of the same fish (anchor and positive) and a third training image of a different fish (negative) and generate 64-dimensional embeddings. Triplet loss compares the anchor to the positive and negative inputs and during training maximizes the Euclidean distance between the anchor and negative inputs and minimizes the distance between the anchor and positive inputs.

**C.** Validation loss versus epoch for training shown in **Figure 2C**. Loss decreases and ultimately converges over training. Here, the network is trained for 62 epochs, which takes approximately 20 minutes.

**Segmentation and counting of C. elegans**

To segment and quantify the number of C. elegans and their swarming behavior, we utilized a U-shaped fully convolutional network (Ronneberger et al., 2015), widely recognized as U-net, to create binary identification masks of both worms and swarms. To supervise the training, we created a manually labeled dataset composed of 655 manually labeled, 256x256 sized images, which was then augmented to 13100 labeled images using rotation. We choose a U-Net architecture (Buda et al., 2019) that is publicly available from the PyTorch library. To make the network output per-organism and per-swarm masks, we set the output channel numbers to two and initialized the network via a uniform random distribution of weights bounded by the square root of input features of each convolution layer. During training, we input image segments containing either individual C. elegans, swarms, or both, with the output being the associated segmentation masks. The network was trained with the Adam optimizer using a batch size of 12, a learning rate of $1 \times 10^{-4}$ for 200 epochs. Once trained, we input the network with $512 \times 512$ sized tiled images from our MCAM-96 video sequences, cropped from the stitched gigapixel images with a 25% overlap (i.e., a 384-pixel step size). After the network predicted the segmentation masks for each image, we stitched all the segmentation masks together to form the segmentation results for the original gigapixel images. Once segmented, we used a straightforward pixel-connectivity-based method to count the number of objects (worms or swarms) in each gigapixel segmentation mask (Haralock et al., 1991).

***Depth Tracking***

The multi-camera array microscope (MCAM) imaging system uses multiple high-resolution cameras to simultaneously image across a large field of view (FOV). Although these cameras could be configured such that there is no spatial overlap between them, or a limited amount of spatial overlap that just enables effective image stitching, the system presented in this work utilized a larger degree of overlap to enable depth detection (see **Supplementary Figure 2A-B**). Specifically, the cameras used rectangular sensors (2432 x 4320 pixels; 5:9 form factor) and were configured such that the FOV of horizontally adjacent cameras overlapped by slightly more than 50% and the FOV of vertically adjacent cameras overlapped by approximately 10% to enable accurate stitching along the vertical direction. Therefore, any object within the collective FOV of the MCAM was captured by at least 2 cameras, with the exception of the cameras along the vertical edges (at the end and start of horizontal rows (see **Supplementary Figure 7B**).

The fact that adjacent cameras have overlapping fields of view means that, for a large central subset of each frame, there exists stereoscopic image data. Stereoscopic imaging is a robust method of estimating depth information from two 2D images (Birchfield et al., 1999). In essence, stereo-depth estimation process utilizes the difference in how objects are perceived across the two image viewpoints to estimate an object's distance along the optical axis at one or more points, much in the same way that the relative lateral disparity between objects seen by our own two eyes can be used as an indication of object depth.

**Depth-from-stereo methodology**

The key to establishing a depth estimate from a stereo pair of images is mapping pixel disparity of a particular point on the sample of interest to a physical distance. Supplementary Figure 7D shows the geometry of a camera pair from the setup used in this work. The goal is to move from the total pixel disparity on the captured images ($d = d_1 + d_2$) to a physical object depth (Z). Under the assumption that our lenses and sensors obey certain geometric constraints (Arellano et al., 2016), we can use a simple similar triangles relationship (Eq. 1) to establish the mapping between pixels and physical geometry:

$$\frac{\mathrm{d}}{\mathrm{L}} = \frac{\mathrm{B}}{\mathrm{Z}} \quad \rightarrow \quad \mathrm{Z} = \mathrm{BL}/\mathrm{d} \qquad \text{Eq. (1)}$$

This relationship is sketched in **Supplementary Figure 7D**. Here, B is the inter-micro-camera spacing (19 mm) and L the lens-to-sensor (i.e., image) distance (~29 mm), which is computed via the lens-maker equation for f=25 mm lenses assuming a 150 mm working distance. We assume both parameters are fixed, as they are based on the mechanical construction of the MCAM system and remain constant across all camera pairs. Relevant parameters are listed in **Supplementary Figure 7E**. The final pixel disparity parameter of interest, d, which is the sum of the feature offset distances from the centers of each image pair measured along each image plane, is a function of object depth and dynamically changes based on the relative positioning of a point of interest across each pair of adjacent micro-cameras. To estimate object depth (Z) from our stereo system, we thus need a way of estimating the pixel disparity d on a per-object basis.

**Depth-from-stereo computational pipeline**

The first step to depth estimation is thus to select corresponding regions of interest (ROI) from images of adjacent camera pairs, which allows us to isolate a single object (e.g., the tail of a particular zebrafish). Our process of ROI selection utilizes an object detection algorithm, as detailed in the description of the Object Detection Pipeline (Supplementary Figure 5). However, any method of interest can be applied to isolate specimen areas of interest. After selecting an object or specimen area of interest, we then apply a feature detection pipeline to estimate the average pixel disparity between the two perspectives of the single object of interest. This process is illustrated in **Supplementary Figure 7F**.

There are two critical components of depth estimation from feature detection: 1) feature creation (keypoint detection and description), and 2) feature matching. When combined, these two components allow us to find pairs of keypoints within stereoscopic images that are likely to represent the same physical location on the object. To detect and describe keypoints, we utilized the OpenCV implementation of ORB (Oriented FAST and Rotated BRIEF) (Rublee et al., 2011). The ORB process first performs keypoint detection on each sub-image, attempting to find areas which contain high amounts of unique information, which is performed using a modified version of the FAST (Features from Accelerated Segment Test) algorithm (Rosten et al., 2006). Once keypoints are identified, the BRIEF (Calonder et al., 2010) (Binary Robust Independent Elementary Features) algorithm is used to form a feature descriptor. In this case, the descriptor is a binary feature vector that summarizes the spatial contents of a fixed-sized patch of pixels centered at the keypoint location.

After generating a list of candidate features and corresponding feature vectors for each of the two images comprising the stereo-image pair, we then match keypoints between images by minimizing the Hamming distance of their feature descriptors vectors (with all combinations of features tested via brute force). We apply a maximum Hamming distance threshold to the matching process to exclude spurious or poor matches from being included. To further increase the quality of keypoint correspondence, we applied a geometrically constrained RANSAC (Fischler et al., 1981) to the matching keypoint pairs. This last step ensured that the final set of matched keypoints conforms to a Euclidean transform, which is a constraint driven by the mechanical properties of our imaging system. Finally, we computed the average distance between matched keypoints (in pixels), and multiplied this by the pixel size ($p = 1.4$ µm) to obtain an estimate of the object's pixel disparity, d. As an average, this statistic exhibited sub-pixel-level resolution. Using Eq. 1, combined with the known physical parameters of the MCAM (b and L) we transform our digital measurement of d into a physical estimate of the depth Z.

**Depth-from-stereo experimental verification**

Once we established our depth detection pipeline, we constructed an experiment to validate the MCAM's depth detection accuracy and range. For this experiment, a standard 1951 USAF resolution test chart was placed on a motorized stage within the imaging area of the microscope. The stage used had a calibrated accuracy of 2 µm in the direction of travel (vertical) across all cameras and extending well beyond (5 mm+) the expected lens depth of field (approx. 0.5 mm).

For validation, the stage was moved between 50 positions, each 100 µm apart, across a total depth range of 5 mm. At each position, a set of images was captured (one from each of the two utilized micro-cameras). Since the resolution target (shown in **Supplementary Figure 7G** includes few but accurate features, the standard ORB keypoint detection and feature description process was replaced with a simple centroid detection step. The remaining steps in the depth detection process remain unchanged. By removing the need for feature detection, we isolate the validity of tracking depth with the MCAM imaging system from the ability of algorithms to detect features accurately and robustly, which can vary from sample to sample.

After applying centroid detection on 3 bars within the resolution target, the relative position of each centroid is compared across adjacent cameras to estimate depth, following Eq. 1 and using the parameters found in Fig. **Supplementary Figure 7E**. The results of this experiment are shown in **Supplementary Figure 7H**, which show a clear match between predicted and measured depth across the entire 5 mm region, which extends well beyond the depth of field of the MCAM (approximately 0.5 mm).

The axial resolution of MCAM depth measurement can be directly related to the size of the image sensor pixel. As noted above, if multiple features are available per object area of interest, then statistical averaging can yield "sub-pixel" resolvability. However, assuming just one feature is used, we can estimate this resolution using Eq. 1. Due to the inverse relationship between sensor disparity d and Z, the effect of a single pixel difference changes depending on total disparity. To obtain an approximate value of depth precision, we use the effect of a single pixel change when the sample is at a depth of 150 (the working distance of the MCAM system). Thus, plugging in the image distance L = 28.9 mm, the baseline distance b = 19 mm and $Z_a$ = 15 cm into Eq. 1 we obtain $d_a$ = 3.66 mm. We can then use this quantity to find the effects of a 1 pixel shift on Z by computing $Z_b = bL/( d_a + p )$ = 149.943 mm, where p = 1.4 µm is the sensor pixel size. The result suggests that that for every pixel of displacement measured on the sensor, the object of interest is axially displaced by approximately $Z_a - Z_b$ = 57 µm in the sample plane, which can be used as an approximate measure of MCAM depth resolution via stereo-matching.

**Supplementary Figure 7**

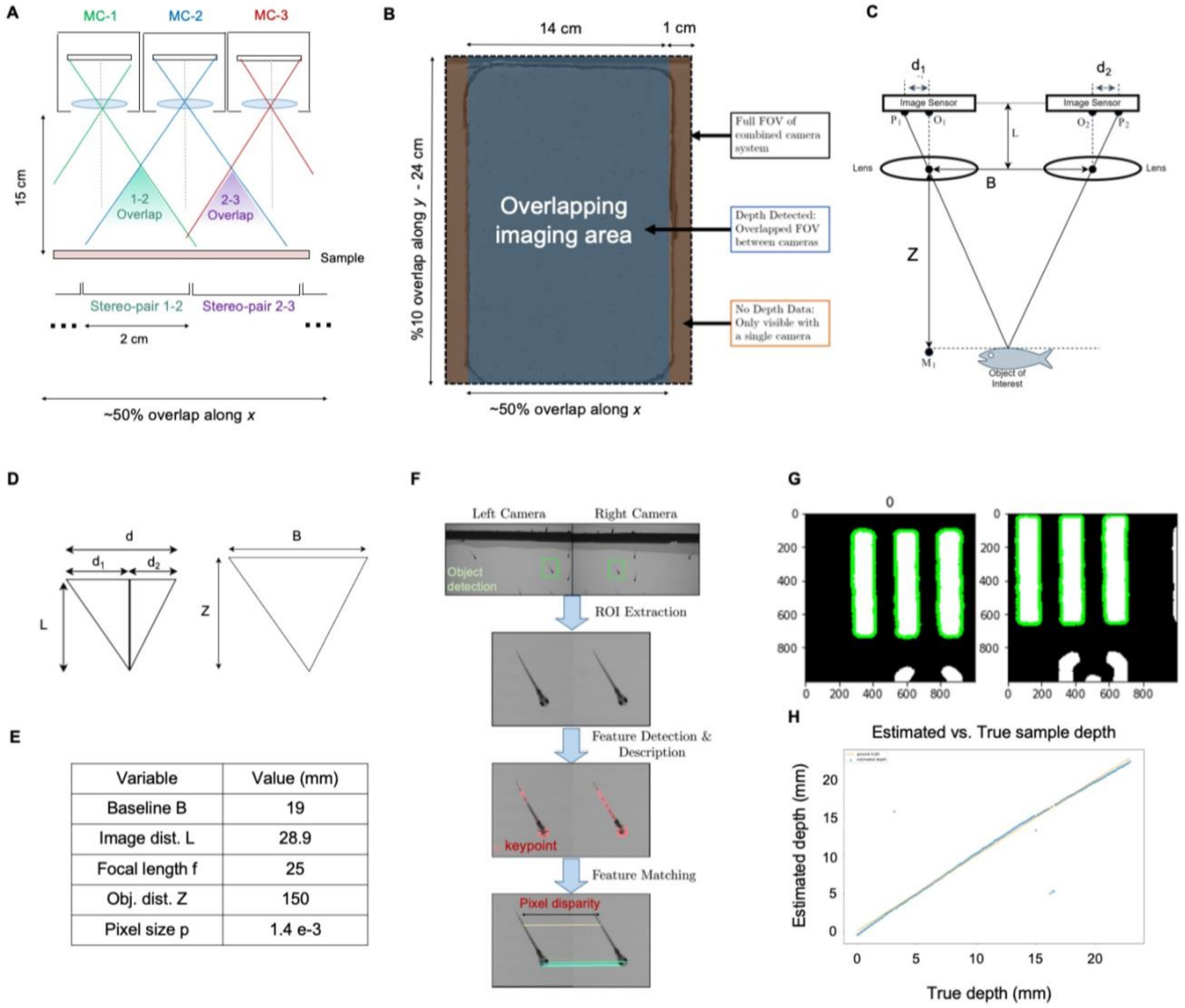


| Variable | Value (mm) |
|---|---|
| Baseline B | 19 |
| Image dist. L | 28.9 |
| Focal length f | 25 |
| Obj. dist. Z | 150 |
| Pixel size p | 1.4 e-3 |

**Supplementary Figure 7 | MCAM Stereoscopic imaging**

**A.** Diagram of MCAM imaging geometry for stereoscopic depth tracking with marked stereo-pair regions

**B.** Stereo-pair regions exist within an overlapping imaging area (other than a small boarder region).

**C.** Schematic of stereoscopic imaging arrangement with variables of interest marked. Object distance (Z) is calculated via trigonometry, after digital estimation of $d_1$ and $d_2$.

**D.** (d) Similar triangles from (c) used for depth estimation via Eq. 1.

**E.** Table of variables and typical experimental values.

**F.** Feature Detection and Matching Pipeline. Matching ROI extracted from a stereo pair, a feature description algorithm is applied to each image and are then matched to compute $d_1$ and $d_2$ for pixel disparity measurement

**G, H.** Experimental results, indicating approximately 100 µm resolution over the entire depth range. Outlying points are due to the inability to accurately match image pair features.

***Fluorescent organism detection and tracking pipeline***

1. Fluorescence Imaging Hardware
To enable the MCAM-24 setup to capture epi-fluorescence images and video, we made two simple hardware modifications:

1) We added two excitation LEDs positioned on either side of the array (each with a 500 nm short pass filter) in an angled epi-illumination geometry, as shown in **Supplementary Figure 8A**.

2) We included a 6x4 array emission filter, arranged in a customized filter holder, placed directly in front of the micro-camera array optics. For single-channel fluorescence measurements, we used a 6x4 array of Chroma 510±10.0nm filters (12 mm square) emission filters **Supplementary Figure 8B**. As outlined in the main text, the MCAM is also capable of recording dual-channel fluorescence video. To achieve this functionality, we used an interleaved combination of Chroma 510±10.0nm filters for green fluorescent *Tg(elavl3:GCaMP6s)*, and Chroma 610±37.5nm filters for red fluorescent *Tg(slc17a6b:loxP-DsRed-loxP-GFP)* signal acquisition. We used 12 of each filter **Supplementary Figure 8E**. The two filter types were interleaved in columns to utilize the MCAM's 50% overlap in the horizontal direction, as shown in **Supplementary Figure 8F**.

2. Data Pre-processing
Before recording both single and dual-channel fluorescent video, we first captured one snapshot MCAM image in complete darkness to obtain a measurement of fixed-pattern sensor noise (i.e., hot pixels). We additionally imaged once with the excitation sources on and with the swim arena in place, but without any organisms within the arena, to obtain an arena-specific background image. Subsequently, we captured video of freely swimming fish (approx. 150 seconds per experiment on average), saving the raw image steam as a sequence of (.bmp) file sets (24X images per set).

Stitching composite images for fluorescent organism experiments is more challenging than with non-fluorescent image data, as most of part of the captured image is black (i.e., non-fluorescent background) and there are few features to assist with accurate inter-camera alignment. Accordingly, we used existing stitching software (PTGui) that facilitates image stitching via a pre-captured template. To generate the template, we placed a patterned target directly beneath the same transparent swim arena filled with the same amount of water (as the water produces refractive distortions that must be accounted for). We imaged this patterned target under matching imaging conditions with all cameras in the MCAM-24 and stitched them with PTGui software, saving the associated template. For dual-channel fluorescent imaging experiments, we stitched each fluorescent channel separately to generate two templates. Finally, we were able to apply the template(s) to stitch subsequently captured video frames.

3. Localizing and Tracking
After each image set was stitched to form a video of stitched composite frames, we next used OpenCV to segment and localize the fish for each individual frame via a simple thresholding algorithm. This process was performed for each fluorescence channel separately for dual-channel data (see **Supplementary Figure 8G**). We calculated the center point of each segmentation mask and used two consecutive center points to obtain the Euclidean distance, which was then used to calculate speed, as specified in detail below (see **Supplementary Figure 8H**). For instances in which the fish moved too fast to track, frames were not used for analysis; however, omitted frames were accounted for in final speed calculation.

Once the center point of each segmentation mask for each individual organism was localized, we used each organism's 2D x-y coordinates $(x_i^t, y_i^t)$ for organism $i = 1,2,..M$ at frame t to predict organism speed and trajectory. We accomplished this via an optimization approach that allowed us to jointly identify matching organisms and their corresponding 2D spatial displacement in subsequently recorded video frames. We aimed to minimize the total cross-organism displacement D by finding the binary assignment variables $g_{i,j}^t$ that minimize the sum of per-organism displacements g between two consecutive frames,

$$D = \sum_{i,j}^{M} g_{i,j}^t d_{i,j}^t$$

Here, the per-organism displacement $d_{i,j}^t = \sqrt{\left(x_i^t - x_j^{t-1}\right)^2 + \left(y_i^t - y_j^{t-1}\right)^2}$ is the displacement if the organism at $(x_j^{t-1}, y_j^{t-1})$ in frame $t-1$ travels to the position $(x_i^t, y_i^t)$ in frame t. Since we assume that only one organism can travel to one specific position within a particular next frame, we assume $g_{i,j}$ must satisfy,

$\sum_i^M g_{i,j} = 1, \sum_j^M g_{i,j} = 1, g_{i,j} \in \{0,1\}$.

In other words, if we construct a matrix whose $i, j^{th}$ element is $g_{i,j}$, the only one element of each column or row of such a matrix can have the value 1, and the remaining elements of the matrix must be zero. $g_{i,j}^t$ = 1 if the $j^{th}$ fish at frame $t-1$ swims to $i^{th}$ position at frame t, and is $= 0$ otherwise. We attempt to find the assignment of binary values within the matrix g that minimize the total travel displacement with the Hungarian algorithm, which has a time complexity of $O(n^3)$. Once we figure out the binary assignments of g, we can calculate the distance that each fish travels between frame t and $t-1$. For example, if it is determined that $g_{3,5} = 1$, this implies that zebrafish 3 at frame t moves from position 5, $(x_5^{t-1}, y_5^{t-1})$ at frame $t-1$. Hence, we can compute the total displacement of zebrafish 3 between the two frames as $d_{3,5}^t = \sqrt{(x_3^t - x_5^{t-1})^2 + (y_3^t - y_5^{t-1})^2}$, and its approximate speed is $d_{3,5}^t/e$, where e is inter-frame time delay.

3. Quantitative Fluorescence Analysis

Captured images were first de-Bayered into three channels (red, green, and blue). We used only the green channel for GFP analysis from micro-cameras with the associated GFP filter (in both single and dual imaging). For RFP analysis in dual-channel imaging, we used only the red channel from micro-cameras with the associated RFP filter. We computed the total energy within each fluorescent channel per organism per frame by computing the sum of associate GFP/RFP channel intensity values within each segmented fish area **Supplementary Figure 8G**. For accurate comparison in the dual-channel fluorescent imaging scenario, we required that the same number of pixels are considered for both channels, which we achieved by first registering the two segmentation masks from the green and red channels and then producing a common segmentation mask as the union of these two masks. Using the common mask, we then measured each channel's total energy.

**Supplementary Figure 8**

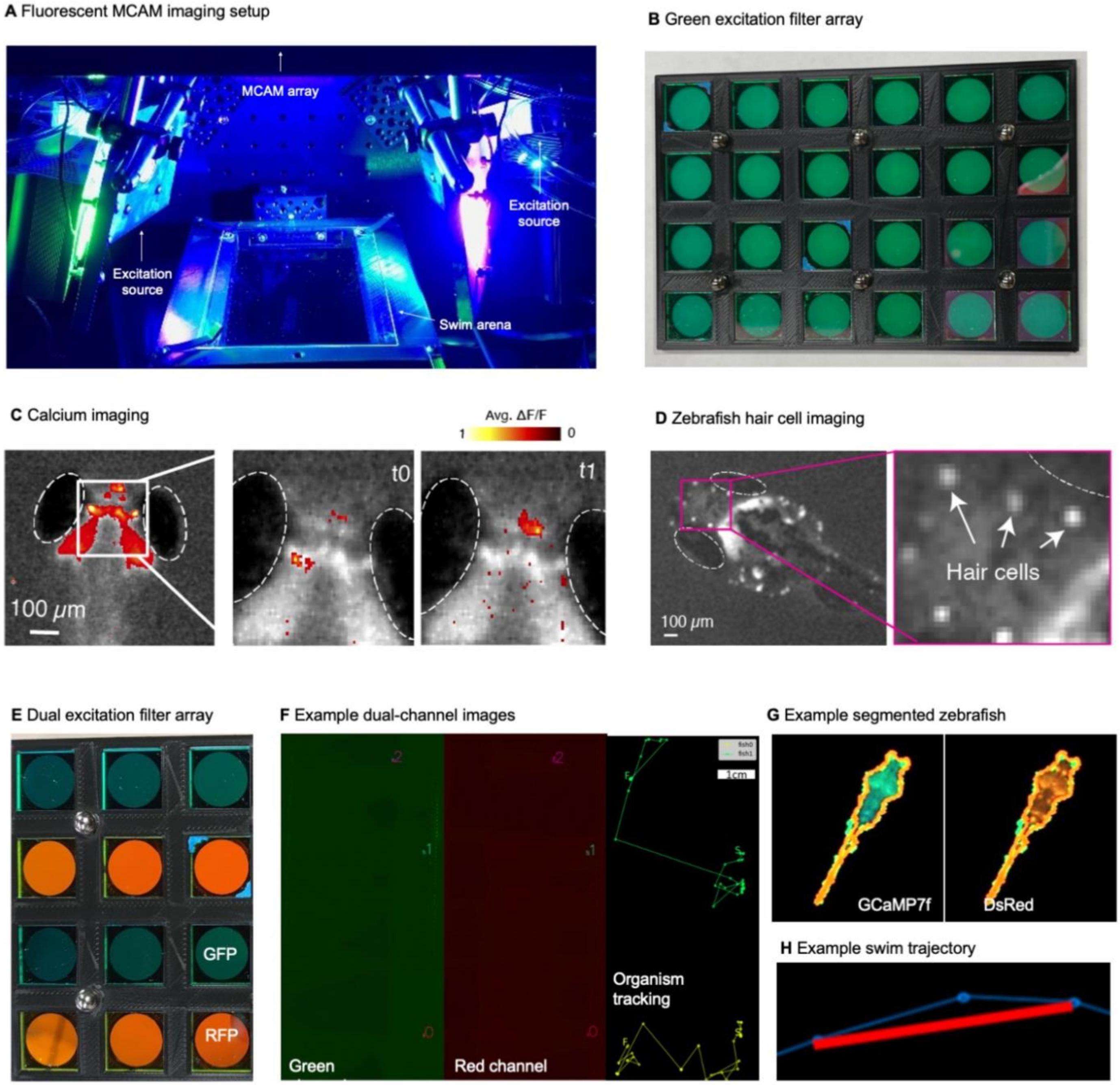

**Supplementary Figure 8 | MCAM Fluorescence Imaging.**

**A.** Fluorescent MCAM imaging setup with two excitation LEDs at an angle above arena in epi-illumination configuration.

**B.** 24 emission filters within array for MCAM imaging of green fluorescent proteins.

**C.** Snapshots of single sensor fluorescence images (30 Hz) of neural activity in 6 day old *Tg(elavl3:GCaMP6s)* zebrafish embedded in agarose.

**D.** Snapshot and zoom in of 6-day old zebrafish with fluorescently labeled hair cells demonstrates near cellular resolution.

**E.** 12 of 24 emission filters within array used for simultaneous capture of GFP and RFP fluorescence signal for ratiometric video of neural activity within freely moving organisms (12 or 24 shown here).

**F.** Example stitched MCAM frames across 8x12 cm FOV of simultaneously acquired green (GFP) and red (RFP) fluorescence channels, with 3 freely swimming zebrafish larvae exhibiting both green *Tg(elval3:GCaMP6s)* and red *Tg(slc17a6b:loxP-DsRed-loxP-GFP)* fluorescence emission. At right is tracked swim trajectories over 240 sec acquisitions, from which swim speed is computed.

**G.** Example of automated segmentation of a double transgenic fish in the red and green channel from the recording shown in F.

**H.** Demonstration of swim trajectory calculation.

**Supplementary Videos**

**Supplementary Movie 1 | 3D rendering of MCAM**

Three-dimensional rendering of the MCAM hardware. Corresponds to discussion of **Figure 1A**.

**Supplementary Movie 2 | High-resolution, large-FOV larval zebrafish**

Brightfield video of freely swimming, 8-day old zebrafish, recorded at approximately 1 frame per second for 1 hour (2000 frames shown here). Randomly selected zoom-in locations at two spatial scales demonstrate ability to resolve individual organism behavior at high fidelity across full 16x24 cm field-of-view. Corresponds to discussion of **Figure 1D**.

**Supplementary Movie 3 | Example output of gigadetector algorithm**

Brightfield video of freely swimming, 8-day old zebrafish, showing output of automated organism detection software marked as a bounding box and label around each larva. Corresponds to discussion of **Figure 2A**.

**Supplementary Movie 4 | Zebrafish GCaMP based calcium imaging**

Fluorescence calcium imaging of spontaneous neural activity in agarose restrained zebrafish imaged at 10 Hz (scale bar: 10 microns). Corresponds to discussion of **Figure 3D**.

**Supplementary Movie 5 | Dual mode zebrafish fluorescent imaging**

Video of freely swimming transgenic zebrafish expressing the green fluorescent genetically encoded calcium sensor (GCaMP6s) and stable red fluorescent protein (dsRed) in almost all neurons and only GABAergic neurons, respectively. Simultaneous dual-channel MCAM recording of both red fluorescence (dsRed) and green fluorescence (GCaMP6s) allows ratiometric measurements by normalizing the fluctuating green fluorescence reflecting changes in neural activity with the stable signal in the red channel. Two color channels are shown on the left across full MCAM FOV, with tracked zoom-ins of each fish shown to right. Corresponds to discussion of **Figures 3G-H**.

**Supplementary Movie 6 | Wild type C. elegans**

Brightfield video of C. elegans wild type (N2) imaged with MCAM-96. Randomly selected zoom-in locations at two spatial scales demonstrate ability to resolve individual organisms. Corresponds to discussion of **Figure 4A**.

**Supplementary Movie 7 | NYL2629 C. elegans strain**

Brightfield video of C. elegans NYL2629 strain. Randomly selected zoom-in locations at two spatial scales demonstrate ability to resolve individual organisms and jointly observe macroscopic behavioral phenomena, in particular the marked tiling of the dish with periodic swarms of C elegans. Corresponds to discussion of **Figure 4B**.

**Supplementary Movie 8 | C. elegans super-swarming behavior**

Brightfield video of C. elegans unc-3 mutant imaged with MCAM-96. Randomly selected zoom-in locations at two spatial scales demonstrate ability to resolve individual organisms and jointly observe macroscopic super-swarming behavioral phenomena. Corresponds to **Figure 4C**.

**Supplementary Movie 9 | Carpenter ant behavior under multiple light/dark cycles**

Brightfield imaging of collective Carpenter Ant behavior using MCAM-96, with ambient lighting sequentially turned on and off every two minutes (6 repetitions). Randomly selected zoom-in locations shown at two spatial scales at right. Corresponds to discussion of **Figure 5B**.

**Supplementary Movie 10 | Slime mold maze traversal**

Video of slime mold *Physarum polycephalum* traversing a custom-designed maze from 4 starting locations, imaged in time-lapse mode with the MCAM-96 over the course of 96 hours. Zoom-ins show ability to observe pseudopodia at high resolution during growth and foraging. Corresponds to discussion of **Figure 5A**.

**Supplementary Movie 11 | Drosophila larva fluorescence demonstration**

Fluorescence video of 3 freely moving Drosophila melanogaster larva expressing GFP imaged at 10 Hz (genotype: w; UAS-CD4tdGFP/cyo; 221-Gal4). Corresponds to discussion of **Figure 3B**

**Supplementary Movie 12 | Drosophila adult bright-field imaging demonstration**

Adult Drosophila during spontaneous free movement and interaction across full MCAM-96 FOV (approx. 16 x 24 cm), with randomly selected zoom-in locations at two scales demonstrating ability to monitor macroscopic and microscopic behavioral phenomena. Corresponds to discussion of **Figure 5C**.

**Supplementary Movie 13 | Slime mold petri dish exploration**

Time-lapse movie (image taken every 15 minutes) shows single slime mold growth from center of petri dish over 46 hours. Petri dish is seeded with multiple oatmeal flakes, and you can observe the characteristic large-scale exploratory behavior of the slime mold over time, as well as finer-scale plasmodia structure.

**Supplementary Movie 14 | Slime mold cytoplasmic flow demonstration**

This was recorded running the MCAM in single camera mode at 10Hz. Cytoplasmic flow, and its reversal, within individual plasmodia, is clearly observable.

**Supplemental Material References:**

Birchfield S, Tomasi C. 1999. Depth discontinuities by pixel-to-pixel stereo. International Journal of Computer Vision. 35(3):269-93. doi.org/10.1023/A:1008160311296

Buda M, Saha A, Mazurowski MA. 2019. Association of genomic subtypes of lower-grade gliomas with shape features automatically extracted by a deep learning algorithm. Computers in biology and medicine. Jun 1;109:218-25. doi.org/ 10.1016/j.compbiomed.2019.05.002

Calonder M, Lepetit V, Strecha C, Fua P. 2010. Brief: Binary robust independent elementary features. European conference on computer vision Sep 5 778-792. doi.org/10.1007/978-3-642-15561-1_56

Fischler MA, Bolles RC. 1981. Random sample consensus: a paradigm for model fitting with applications to image analysis and automated cartography. Communications of the ACM. 24(6):381-95. doi.org/10.1145/358669.358692

Haralock RM, Shapiro LG. 1991. Computer and robot vision. Addison-Wesley Longman Publishing Co., Inc.;

Huang J, Rathod V, Sun C, Zhu M, Korattikara A, Fathi A, Fischer I, Wojna Z, Song Y, Guadarrama S, Murphy K. 2017. Speed/accuracy trade-offs for modern convolutional object detectors. Proceedings of the IEEE conference on computer vision and pattern recognition 7310-7311. doi.org/10.1109/CVPR.2017.351

Kingma DP, Ba, J. 2015. Adam: A Method for Stochastic Optimization**.** Proceedings of the 3rd International Conference on Learning Representations (ICLR 2015).

Krishnamurthy D, Li H, du Rey FB, Cambournac P, Larson AG, Li E, Prakash M. 2020. Scale-free vertical tracking microscopy. Nature Methods. (10):1040-51. doi.org/10.1038/s41592-020-0924-7

Lin TY, Maire M, Belongie S, Hays J, Perona P, Ramanan D, Dollár P, Zitnick CL. 2014. Microsoft coco: Common objects in context. European conference on computer vision 740-755. doi.org/10.1007/978-3-319-10602-1_48

Maaten, LVD., Hinton, G. 2008. Visualizing data using t-SNE. J. Mach. Learn Res. 9, 2579-2605.

Ou X, Horstmeyer R, Zheng G, Yang C. 2015. High numerical aperture Fourier ptychography: principle, implementation and characterization. Optics express. 23(3):3472-91. doi.org/10.1364/OE.23.003472

Romero-Ferrero F, Bergomi MG, Hinz RC, Heras FJ, de Polavieja GG. 2019 Idtracker. ai: tracking all individuals in small or large collectives of unmarked animals. Nature methods.16(2):179-82. doi.org/10.1038/s41592-018-0295-5

Ronneberger O, Fischer P, Brox T. 2015. U-net: Convolutional networks for biomedical image segmentation. InInternational Conference on Medical image computing and computer-assisted intervention(pp. 234-241). doi.org/10.1007/978-3-319-24574-4_28

Rosten E, Drummond T. 2006. Machine learning for high-speed corner detection. European conference on computer vision 430-443. doi.org/10.1007/11744023_34

Rublee E, Rabaud V, Konolige K, Bradski G. ORB. 2011. An efficient alternative to SIFT or SURF. 2011 International conference on computer vision 2564-2571. doi.org/10.1109/ICCV.2011.6126544

Schroff F, Kalenichenko D, Philbin J. 2015. Facenet: A unified embedding for face recognition and clustering. InProceedings of the IEEE conference on computer vision and pattern recognition (pp. 815-823). doi.org/ 10.1109/CVPR.2015.7298682

Singh AP, Schach U, Nüsslein-Volhard C. Proliferation, 2014. dispersal and patterned aggregation of iridophores in the skin prefigure striped colouration of zebrafish. Nature cell biology. 16(6):604-11. doi.org/10.1038/ncb2955

Szeliski R. 2006. Image alignment and stitching: a tutorial, foundations and trends in computer graphics and computer vision. Now Publishers. 2(1):120. doi.org/10.1.1.130.6691

Taigman Y, Yang M, Ranzato MA, Wolf L. 2014. Deepface: Closing the gap to human-level performance in face verification. InProceedings of the IEEE conference on computer vision and pattern recognition (pp. 1701-1708). doi.org/ 10.1109/CVPR.2014.220

Vázquez-Arellano M, Griepentrog HW, Reiser D, Paraforos DS. 2016. 3-D imaging systems for agricultural applications—a review. Sensors. 16(5):618. doi.org/10.3390/s16050618
